## Supplementary Figures for "Brain mitochondrial diversity and network organization predict anxiety-like behavior in mice"

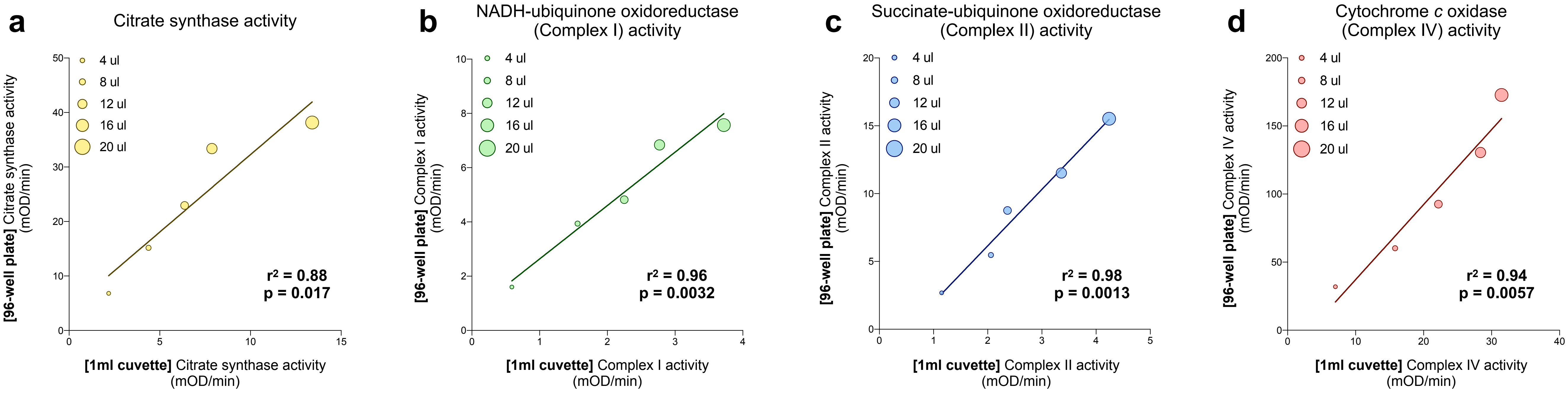


**Supplementary Fig. 1. Validation of miniaturized 96-well plate enzymatic activity assays.** Comparison between miniaturized 96-well plate and traditional 1ml cuvette for enzymatic activities of (**a**) citrate synthase (CS), (**b**) NADH-ubiquinone oxidoreductase (Complex I), (**c**) succinate-ubiquinone oxidoreductase (Complex II), and (**d**) cytochrome *c* oxidase (Complex IV). P values from simple linear regressions. Source data are provided as a Source Data file.

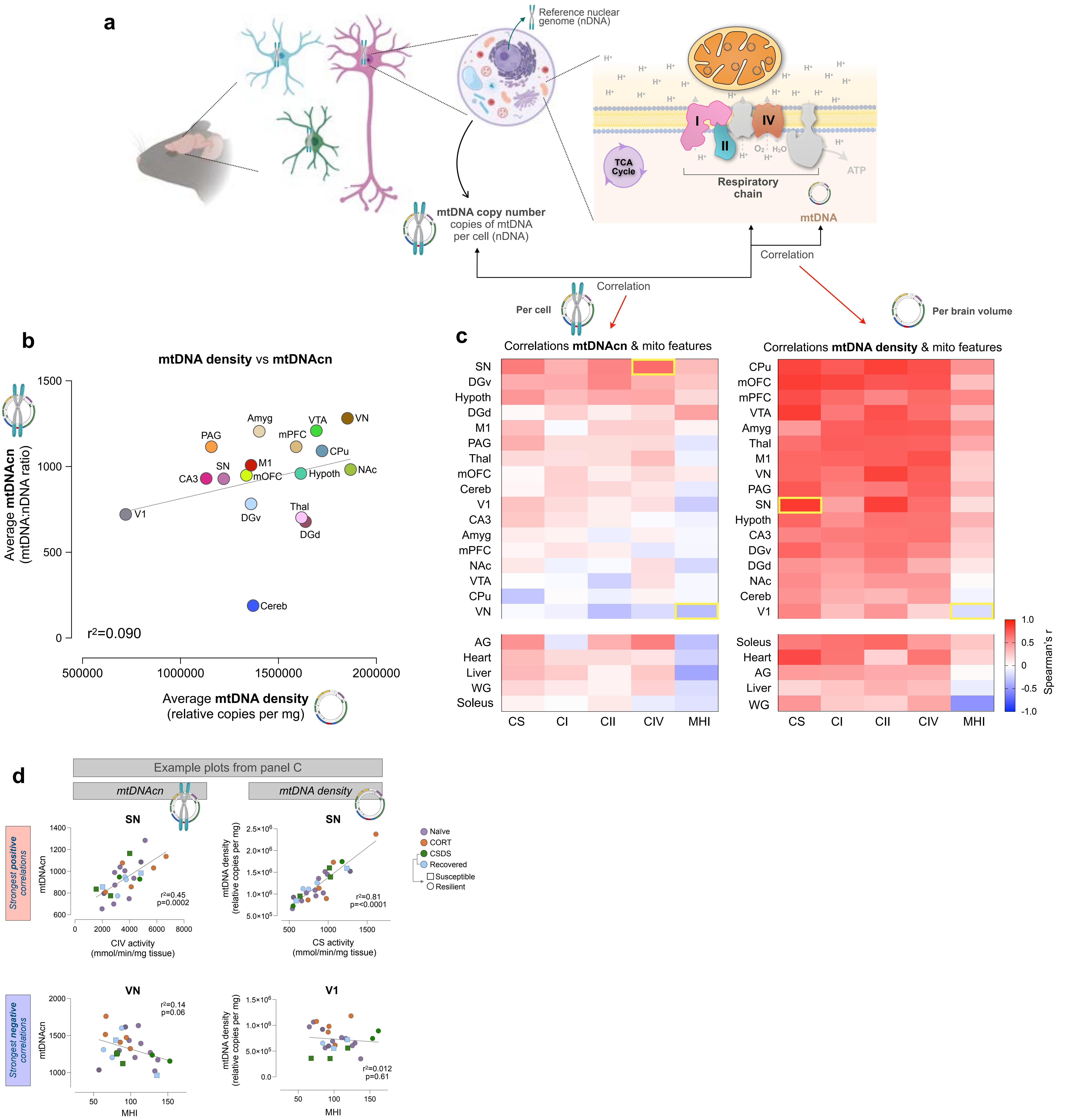


**Supplementary Fig. 2. Comparison of mtDNA copy number versus mtDNA density.** (**a**) Diagram to illustrate the difference between measures of mtDNA copy number (mtDNAcn=mtDNA/nDNA ratio) and mtDNA density (mtDNA copies per mg of tissue). (**b**) Correlation between the average mtDNAcn and average mtDNA density per brain area. (**c**) Correlations between mtDNAcn (left) and mtDNA density (right) with the other five mitochondrial measures in all animals for each brain area and tissue, measured by Spearman's r. The strongest positive and negative brain correlations for each are highlighted with yellow boxes and plotted in (**d**). Two-tailed P values from simple linear regression, upper left: p= 0.0002, upper right: p<0.0001, lower left: p= 0.06, lower right: p=061. Source data are provided as a Source Data file.

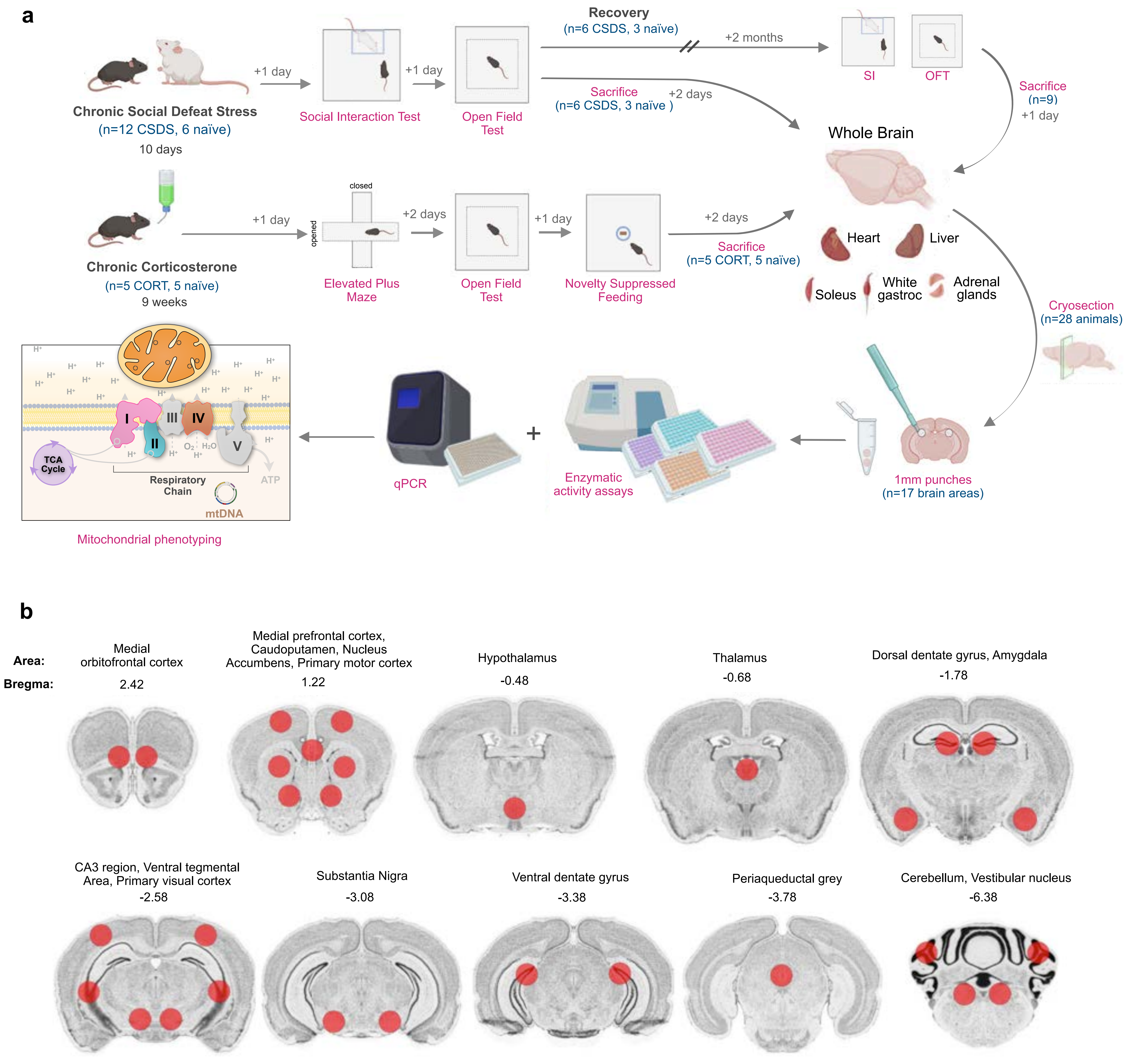


**Supplementary Fig. 3.** (**a**) Experimental design. (**b**) 17 brain areas of interest, labeled by name and distance from bregma, with red circles indicating bilateral punch locations. Images acquired from the Allan Mouse Brain Atlas (Dong, H. W. The Allen reference atlas: A digital color brain atlas of the C57Bl/6J male mouse. *John Wiley & Sons Inc.* (2008)). Abbreviated brain area names are as follows (orders by Bregma coordinates): mOFC, mPFC, CPu, NAc, M1, Hypoth, Thal, DGd, Amyg, CA3, VTA, V1, SN, DGv, PAG, Cereb, VN.

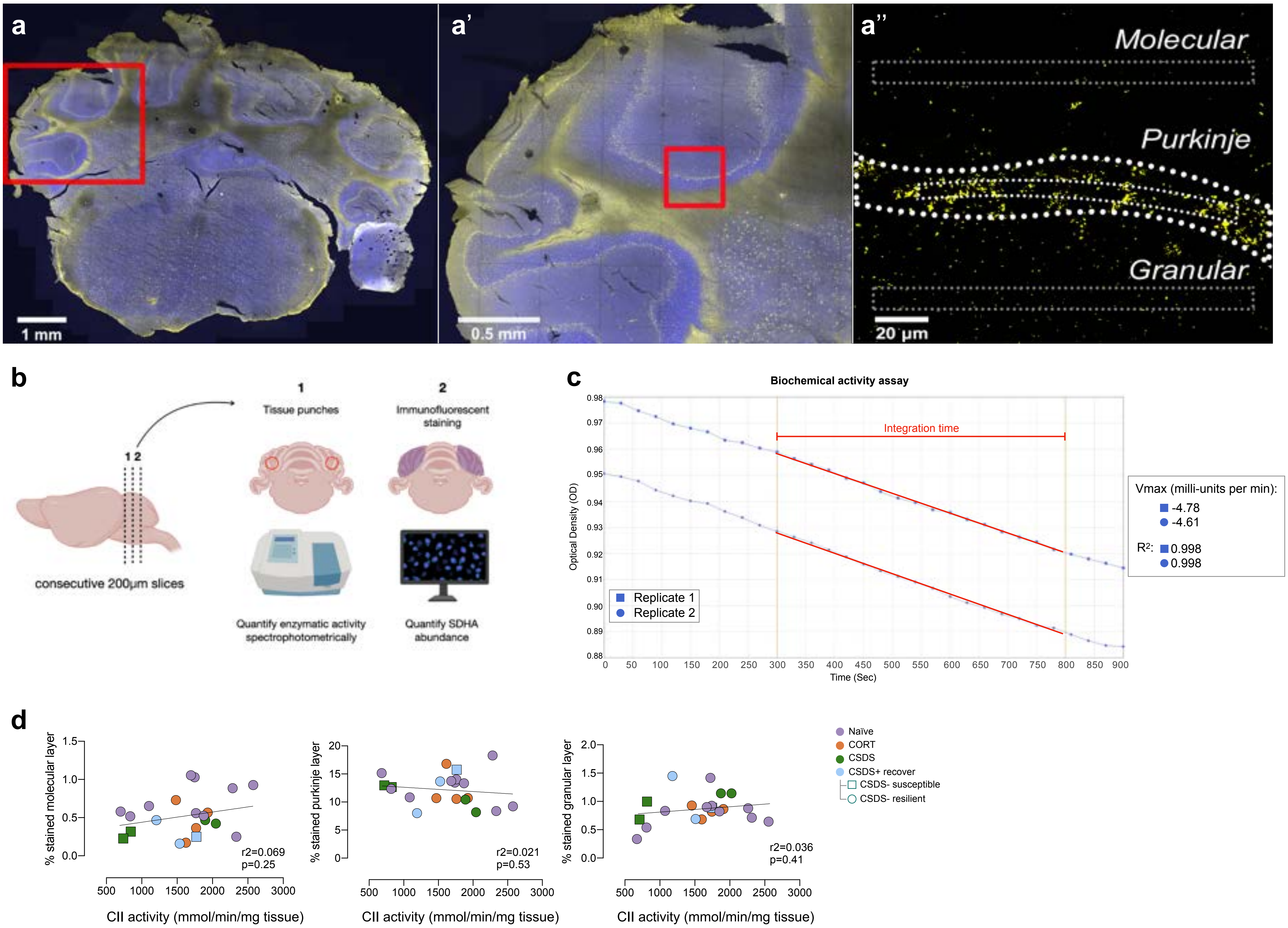


**Supplementary Fig. 4. Cerebellar staining for mitochondrial protein quantification.** (**a**) Coronal 200µm-thick cerebellar slice stained for DAPI (blue, cell nuclei), and the mitochondrial respiratory chain Complex II subunit SDHA (yellow). (**a'**) higher magnification of A, and (**a''**) example high-magnification of SDHA signal along the purkinje cell layer, highlighting three areas of interests used for quantification of SDHA density among the cerebellar molecular, purkinje, and granular layers. Cerebellar staining and quantification was done for each animal from which there were clean slices (n=10 naïve, 4 CORT, 4 CSDS, 3 CSDS + recovery). (**b**) Diagram illustrating that spectrophotometric measurements of CII activity and immunofluorescence confocal microscopy measurements for protein abundance were measured in contiguous 200µm-thick cryosections from the same animals. (**c**) Example raw spectrophotometric trace for CII activity in the cerebellum, integrating OD600 change over 300-800 seconds, showing high agreement between technical replicates. (**d**) Correlations between layer-specific SDHA density (% SDHA-positive pixels per volume) and the biochemical activity of Complex II, demonstrating that protein abundance is not an appropriate surrogate for biochemical activity. P values from simple linear regression. Source data are provided as a Source Data file.

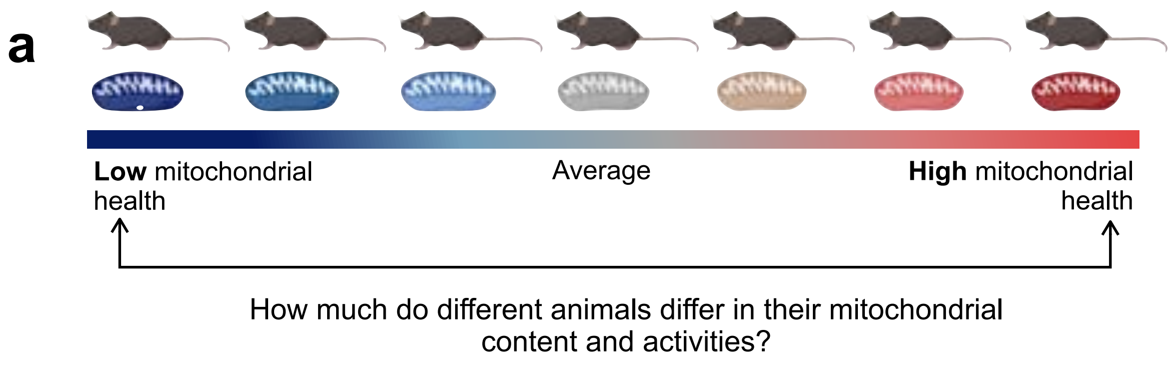


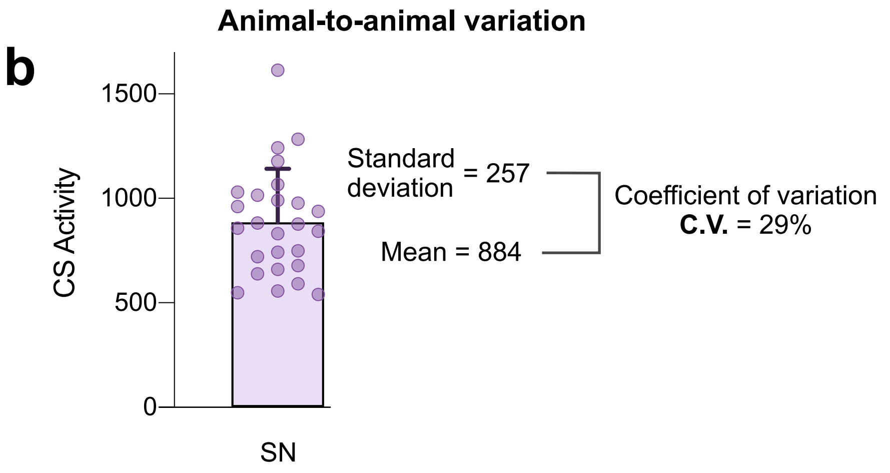


**c**

**Summary inter-individual variation**
(Standard deviation / Mean)

| | CS | CI | CII | CIV | mtDNA | MHI |
|---|---|---|---|---|---|---|
| SN | 29% | 34% | 28% | 34% | 32% | 13% |
| VTA | 27% | 32% | 23% | 29% | 29% | 14% |
| M1 | 23% | 31% | 25% | 30% | 34% | 15% |
| mPFC | 29% | 34% | 26% | 42% | 23% | 16% |
| VN | 34% | 38% | 33% | 33% | 40% | 17% |
| DGv | 23% | 31% | 31% | 30% | 41% | 18% |
| DGd | 26% | 32% | 27% | 38% | 29% | 20% |
| Amyg | 22% | 30% | 36% | 36% | 33% | 21% |
| Thal | 39% | 36% | 30% | 47% | 33% | 22% |
| CPu | 31% | 42% | 33% | 48% | 39% | 23% |
| V1 | 36% | 35% | 32% | 38% | 32% | 25% |
| Cereb | 26% | 35% | 33% | 39% | 37% | 26% |
| Hypoth | 35% | 45% | 44% | 50% | 35% | 28% |
| CA3 | 28% | 39% | 40% | 53% | 45% | 28% |
| mOFC | 41% | 47% | 43% | 58% | 59% | 28% |
| NAc | 30% | 43% | 31% | 39% | 39% | 29% |
| PAG | 42% | 46% | 38% | 60% | 46% | 32% |
| **Brain Avg** | **31%** | **37%** | **33%** | **42%** | **37%** | **22%** |
| AG | 37% | 41% | 32% | 32% | 61% | 21% |
| Liver | 15% | 23% | 14% | 12% | 15% | 8% |
| Soleus | 11% | 22% | 22% | 12% | 23% | 11% |
| Heart | 14% | 20% | 28% | 21% | 17% | 19% |
| WG | 23% | 32% | 34% | 33% | 40% | 22% |
| **Tissues Avg** | **20%** | **27%** | **26%** | **22%** | **31%** | **16%** |
| **Average technical variation** | **4.2%** | **4.4%** | **4.2%** | **8.1%** | **0.6%** | Computed metric |

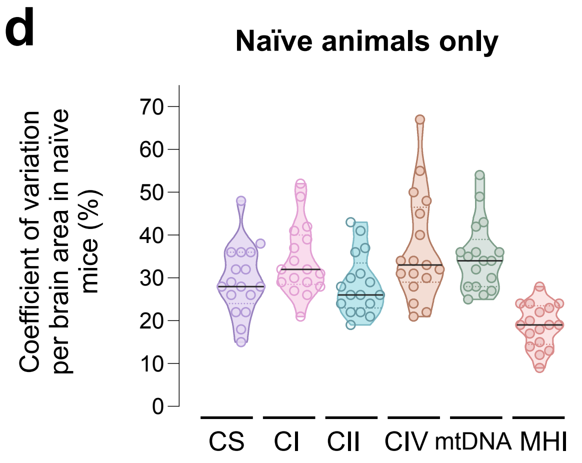


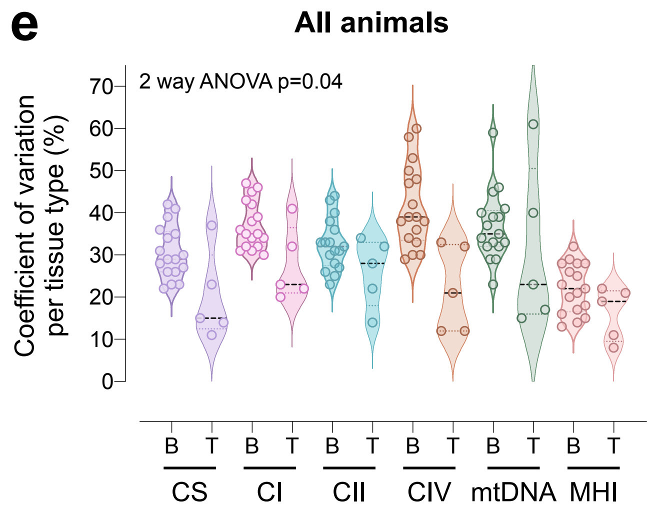


**Supplementary Fig. 5. Animal-to-animal differences in mitochondrial features for each brain area and peripheral tissue.** (**a**) Diagram illustrating the range of mitochondrial content and function across animals. (**b**) Example C.V. (coefficient of variation) calculation for CS activity in the substantia nigra (SN). Each datapoint is an animal (n=27 mice), illustrating the variation in mitochondrial features between animals. (**c**) Individual C.V.s per mitochondrial feature for each brain area and peripheral tissue. (**d**) C.V.s for brain areas by mitochondrial feature for naïve animals only. Each datapoint is the average C.V. for a brain area or tissue (n=22). (**e**) C.V.s for brain areas versus tissues by mitochondrial feature for all animals (naïve, CORT, CSDS), two-way ANOVA, no multiple comparisons. Source data are provided as a Source Data file.

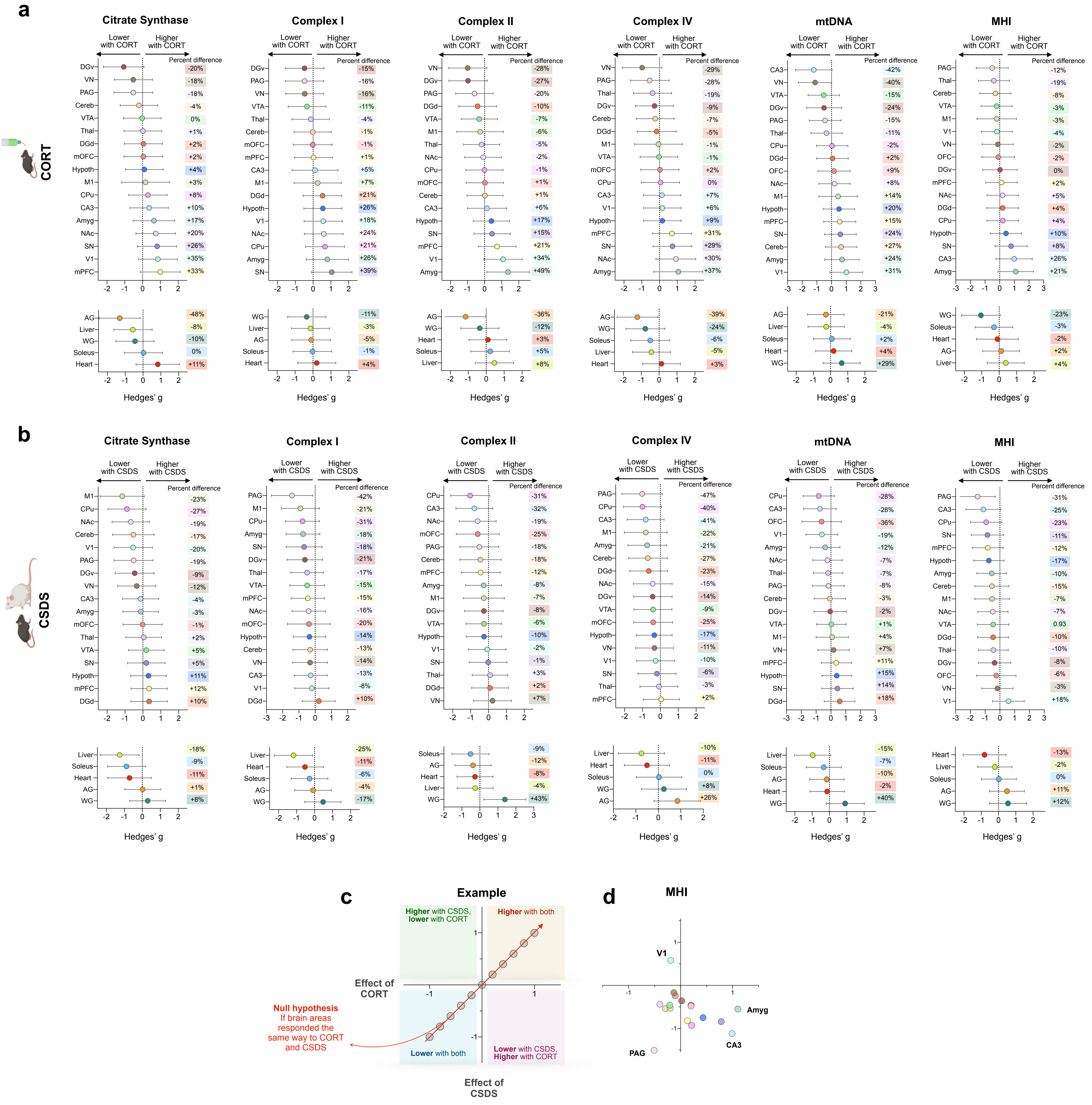


**Supplementary Fig. 6. Effect size of stressors on brain areas' and other tissues' mitochondrial content and functioning.** (**a**) Effect of CORT treatment (n= 5 CORT treated mice and n= 11 naïve mice) and (**b**) effect of CSDS treatment on brain areas and tissues (n= 6 CSDS treated mice and n= 11 naïve mice) compared to naïve mice, quantified by standardized Hedges g with 95% confidence intervals. (**c**) Example plot if both stressors had the same effect on a brain area, with quartiles labeled and (**d**) plot of the effect sizes of both stressors on each brain area's MHI. Source Data are provided as a Source Data file.

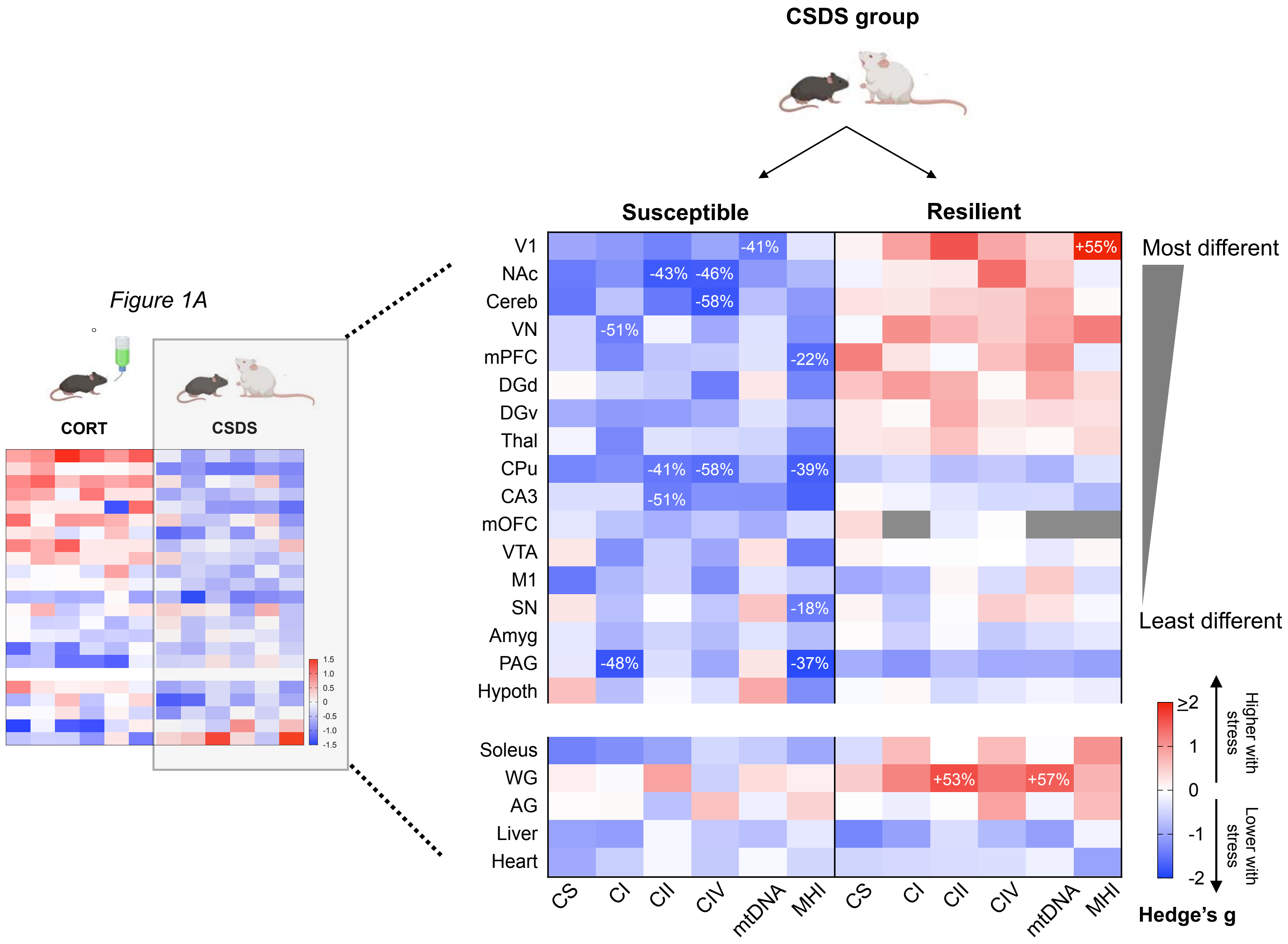


**Supplementary Fig. 7. Exploratory analysis comparing susceptible and resilient subgroups of CSDS mice.** Effect sizes for mitochondrial outcomes shown separately by brain area for animals classified as susceptible or resilient to CSDS (Methods). Effect sizes are Hedge's g, with significant effect sizes (95% confidence interval) labeled with the % change compared to naïve (non-stressed) mice. Source data are provided as a Source Data file.

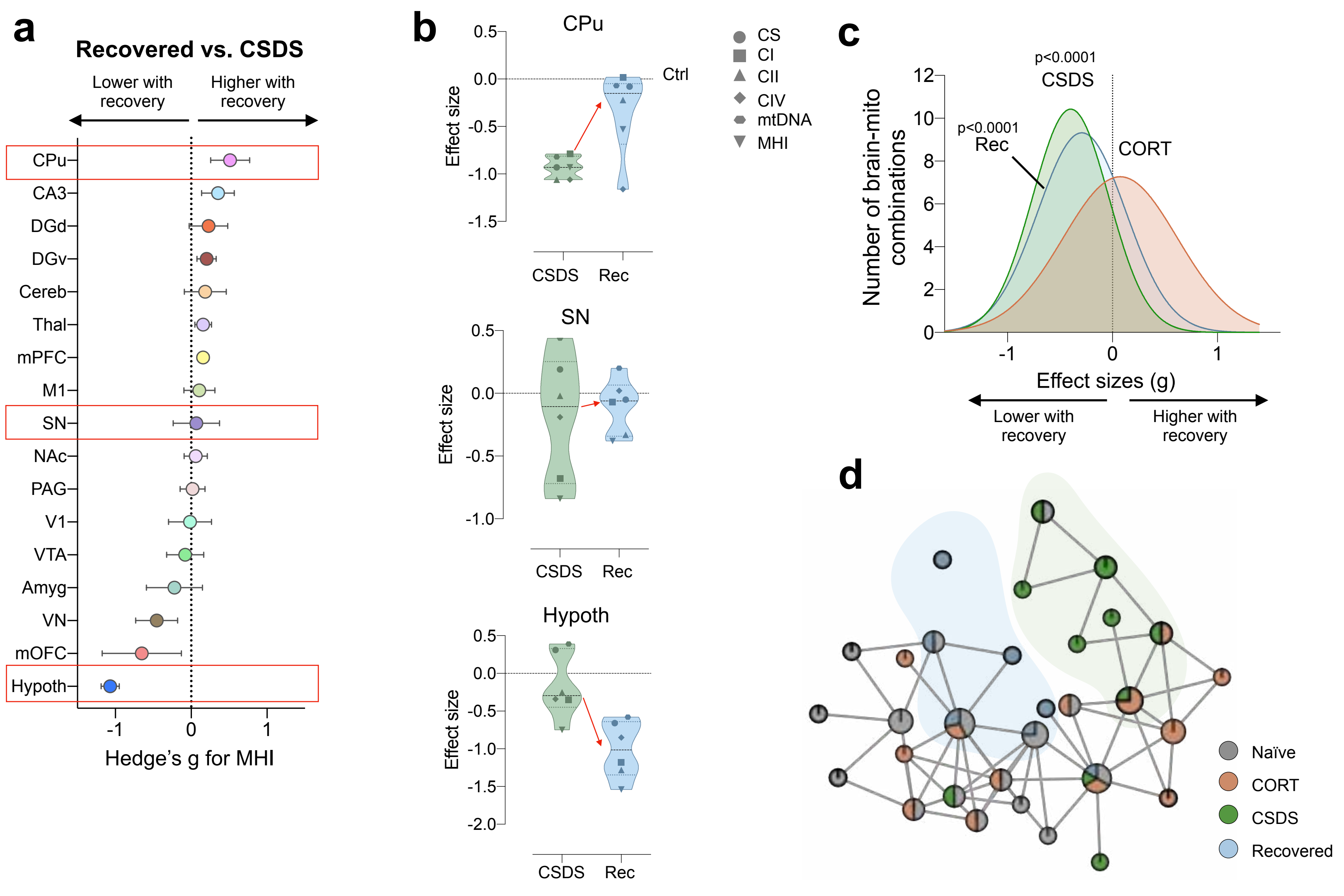


**Supplementary Fig. 8. Effect of recovery from social defeat stress on mitochondrial features across brain areas.** (**a**) Average effect of 2 months recovery (Rec) from CSDS stress across the 6 mitochondrial features for each brain area, as compared to non-recovered CSDS mice, (n= 5 Rec mice, n=6 non-recovered CSDS mice). Effects sizes are quantified by Hedges g, with 95% confidence intervals. MHI in recovered mice was higher than non-recovered CSDS in some areas (Caudoputamen, Hypothalamus-CA3), and lower in others (Hypothalamus, Medial orbitofrontal cortex, Vestibular nucleus). (**b**) Average effect sizes of 6 mitochondrial features of non-recovered CSDS mice and recovered mice compared to naïve mice in three selected brain areas from A (red boxes). (**c**) Frequency distribution of the effect size of CORT, CSDS, and recovery as compared to naïve mice on all 6 mitochondrial measures in all 17 brain areas, gaussian-fitted curve, indicating that recovered mice have lower MHI than naïve mice and marginally higher MHI than CSDS (****p<0.0001, one-sample t test (two tailed)). (**d**) TDA analysis using Mapper of mitochondrial measures in naïve mice, CORT, CSDS, and CSDS-recovered mice, revealing the recovered group to be more similar to the non-recovered CSDS group than to naïve mice. Source data are provided as a Source Data file.

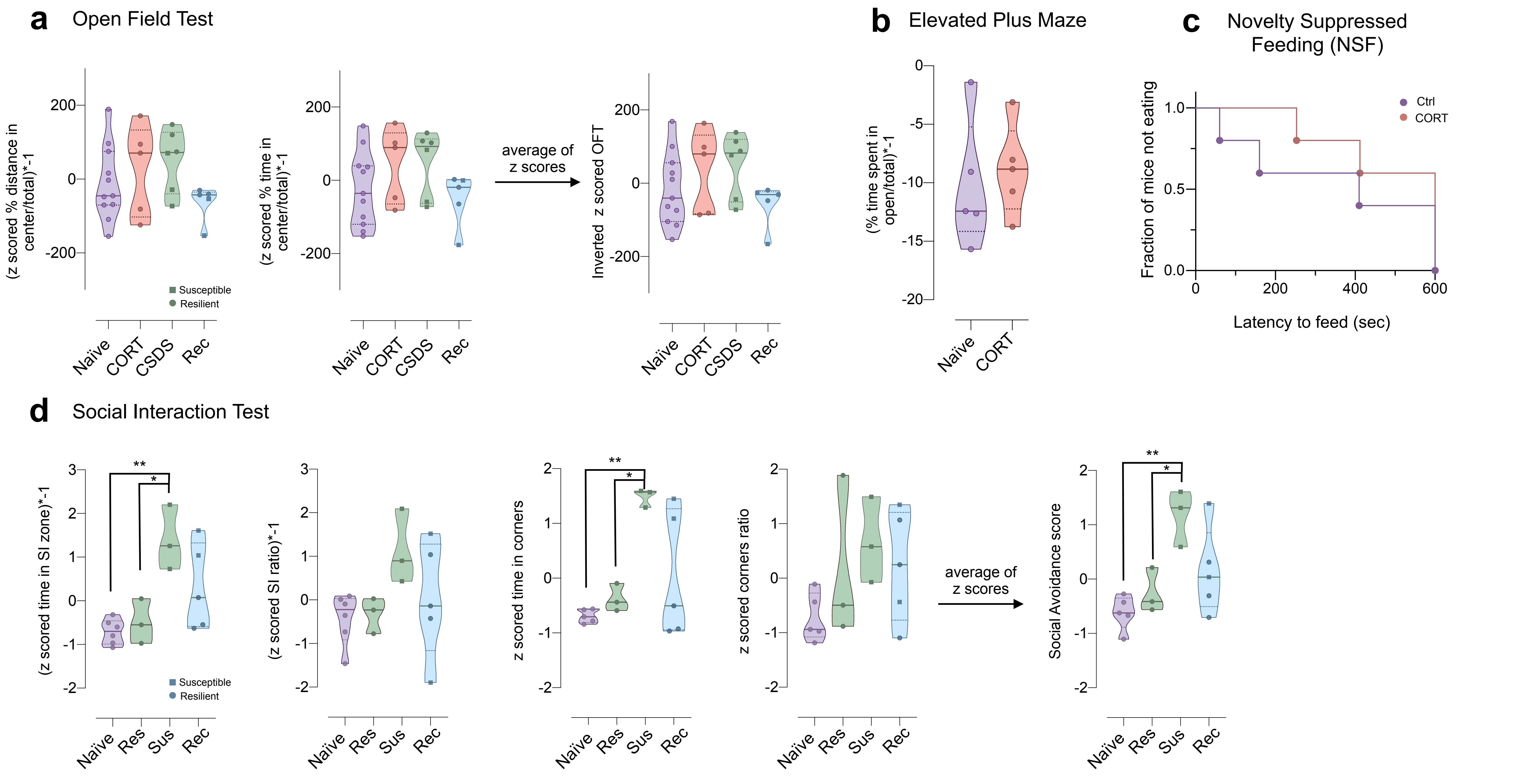


**Supplementary Fig. 9. Behavioral test results** (**a**) To account for both the % time and % distance traveled in the center, the two measures were z-scored and averaged, creating a single Open Field Test (OFT) score. Z-scores were inverted so that a higher OFT score represents less time and distance in the center, indicating higher anxiety. Open field was run for all groups. (**b**) Elevated plus maze (EPM), measuring percent of time spent in open arms/total time, with the score being multiplied by -1 so that higher scores represent less time spent in the open arms, which indicates higher anxiety. EPM was run only on CORT mice. (**c**) Novelty suppressed feeding (NSF) test used to measure the latency to feed in a novel environment, with the test being capped at 600 sec. Survival curve: Mantel-Cox log-rank test. NSF was run only on CORT mice. (**d**) Social Interaction test (SI), represented by a Social Avoidance score for all CSDS mice, separated by susceptible (sus.) and resilient (res.), and recovered (rec). Social avoidance was measured as the average of 4 scores; z scored social interaction ratio, z scored time spent in interaction zone, z scored time spent in corner, and z scored corners ratio. Social interaction ratio and time spent in SI zone were inverted so that higher values for all four measures indicate higher avoidance. z scored social interaction ratio: $p= 0.0035$, $p=0.021$, z scored time spent in corner: $p= 0.0043$, $p=0.026$, Social avoidance score: $p= 0.0045$, $p= 0.040$. Adjust p-values from Tukey's multiple comparison ordinary one-way ANOVA. Source data are provided as a Source Data file.

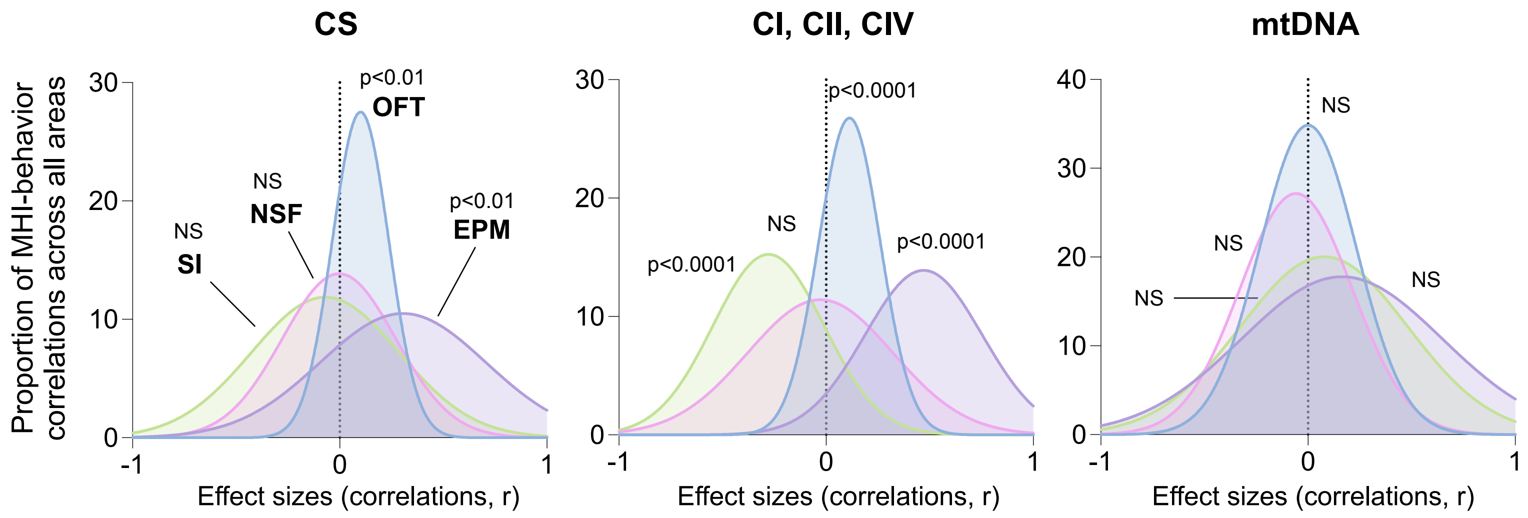


**Supplementary Fig. 10. Frequency distribution of all correlations of mitochondrial-behavior pairs by behavioral test.** Gaussian fits, p values from one-sample t test (two tailed) against the null hypothesis (r=0). CS: OFT p= 0.0019, EPM p=0.0096; CI,CII,CIV: SI, OFT, EPM p<0.0001. Source data are provided as a Source Data file.

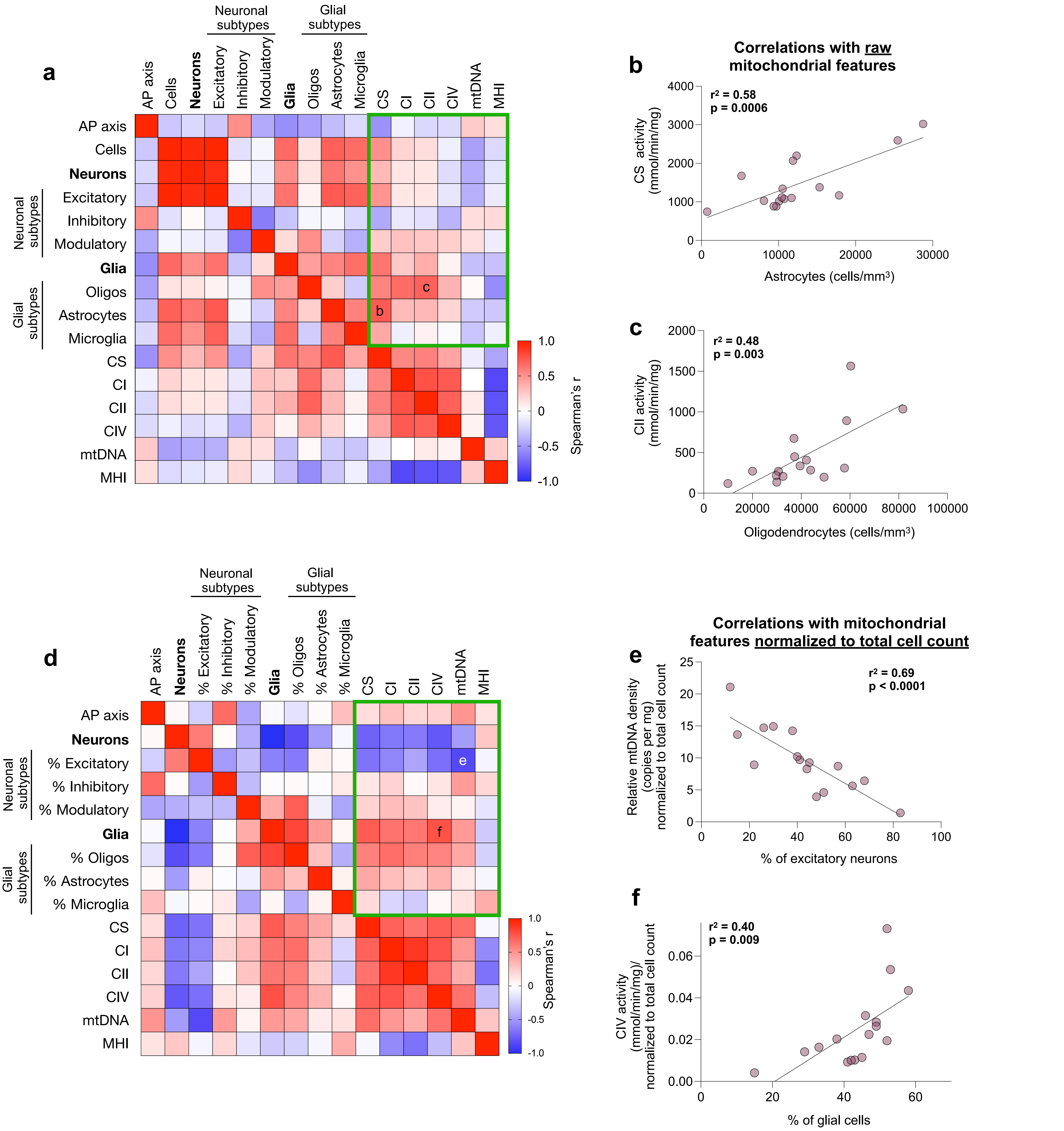


**Supplementary Fig. 11. Correlation of cell type densities and mitochondrial features across brain areas.** (**a**) Data on the abundance or density of various cell types was extracted for each area (n=16 areas, dorsal and ventral not differentiated) from the Blue Brain Cell Atlas[61,62], and correlated with the average (pooled average across all animals) of each mitochondrial feature. The green box indicates the associations between cell type abundances and mitochondrial features. For each brain area, MHI is computed from the average mitochondrial features across all animals. (**b**) Scatterplots for the strongest observed correlations between astrocyte density and citrate synthase activity, and (**c**) oligodendrocyte density (Oligos) and complex II activity. P values from simple linear regression. (**d**) Correlation between the proportion (%) of cell types/subtypes and mitochondrial features normalized to total density of cells for each brain area (i.e., cell count or cellularity). This normalization combines the influence of both mitochondrial content/energy transformation capacity, relative to the number of cell bodies, such that areas with few cell bodies but many mitochondria-rich dendritic arbors have the highest normalized mtDNA density and enzyme activities. (**e, f**) Scatterplots for the strongest correlations from panel d. P values from simple linear regression. Abbreviations: *AP axis*, anterior-posterior axis; *mtDNA*, mtDNA density (relative copies per unit of brain mass). Source data are provided as a Source Data file.

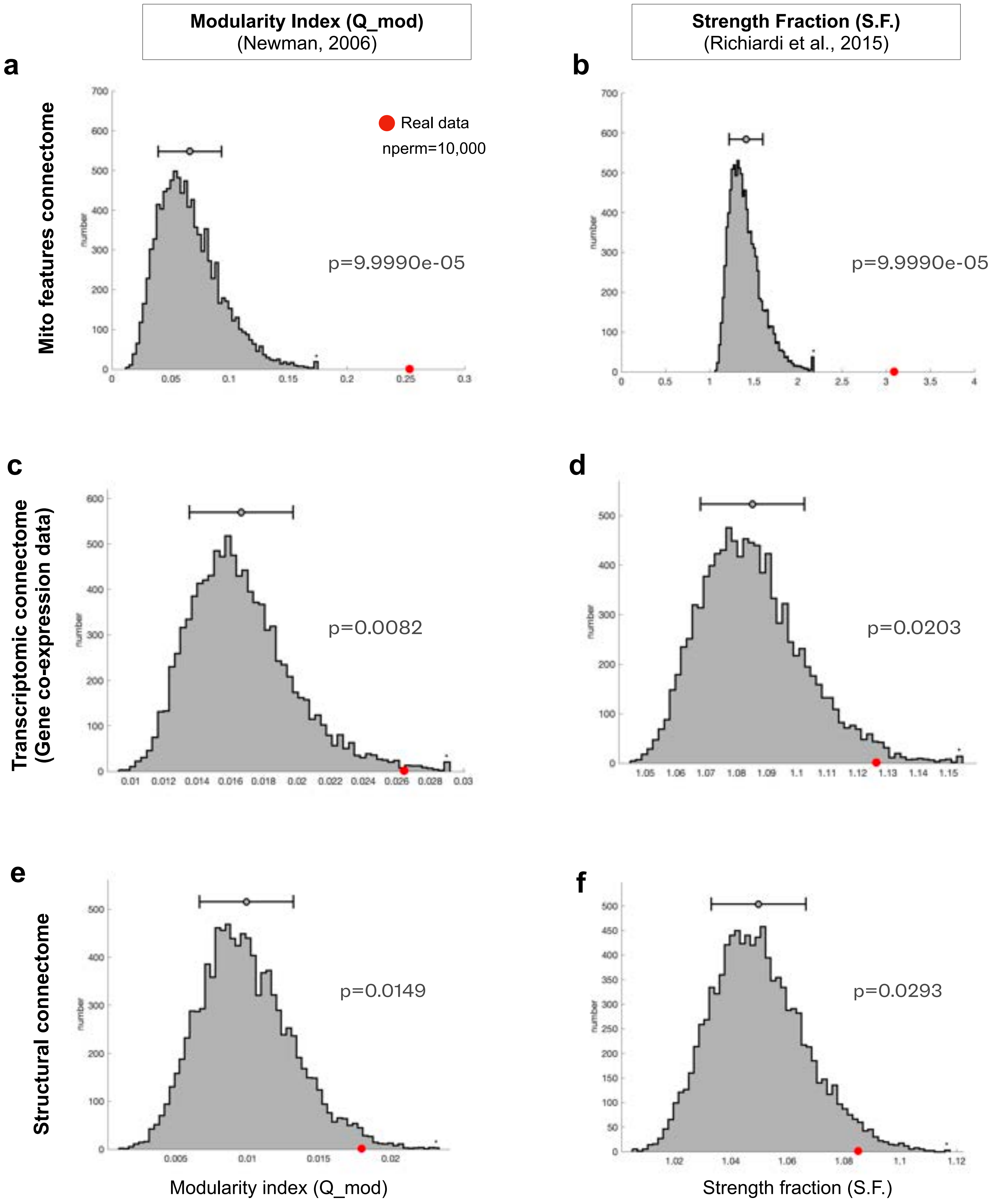


**Supplementary Fig. 12.** To examine whether the functional organization of the brain revealed using mitochondrial features is also evident cross-modally, we utilized two- tailed non-parametric bootstrapping permutation statistics with 10,000 permutations to compare the topology of our mito-derived networks with the expected topology based on the transcriptomic and structural connectomes in the Allen Brain Atlas (see *Online Methods* for details). Specifically, we examined i) whether the tightly knit networks (or communities) observed in the mitochondrial features are more densely connected than expected by chance (**a-b**); and ii) whether the networks derived from mitochondrial features were also more densely connected than expected by chance in other modalities, including gene co-expression data (**c-d**) and EYFP-based structural connectivity (**e-f**). To measure the degree of within-network connectedness we used two established metrics: modularity index (Q_mod; Newman 2006) and strength fraction (S.F.; Richardi et al. 2015). The histograms depict distribution of within-network connectedness generated using 10,000 permutations (or data shuffling), with error bars indicating SEM. The real value of within-network connectedness is shown using red dots across all plots, and we compared whether these real value occur in <0.05% cases. For each modality, and across the two metrics, the networks derived from mitochondrial features were more tightly knit than expected by chance, hence providing convergent multimodal evidence of mitochondrial functional organization overlapping with gene expression and the structural connectome.

**Molecular features of each network**

**Genes fold-difference threshold $Log_2$=1**
OVER: +100% (double)
UNDER: -50% (half)

**Differential expression of biological processes**
FDR=0.05

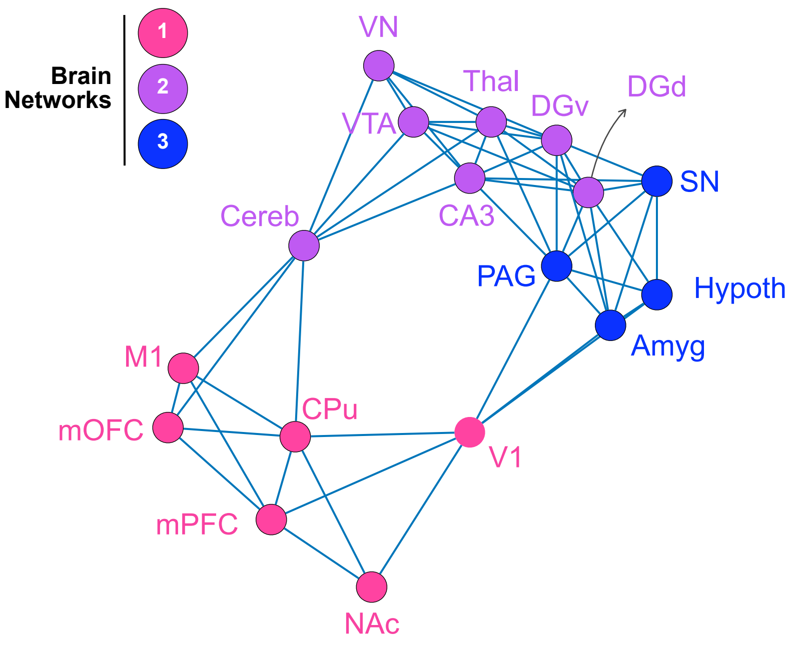


| OVER-expressed — GO biological processes | | |
|---|---|---|
| **1** n=1103 | **2** n=579 | **3** n=799 |
| Synaptic signalling and transmission | Anatomical development and hormonal responses | Calcium regulation |
| Neuronal and dendritic morphogenesis | Ion and calcium membrane transport | Feeding behavior and neuropeptide signaling |
| Enzyme regulation by phosphorylation | Extracellular matrix organization | Metabolic processes |
| Network **1** *vs* **2**/**3** | Network **2** *vs* **1**/**3** | Network **3** *vs* **1**/**2** |

| UNDER-expressed — GO biological processes | | |
|---|---|---|
| **1** n=1130 | **2** n=575 | **3** n=943 |
| Metabolic processes | Cellular movement and locomotion | Cell movement and locomotion |
| Oxygen and chemical sensing | Transynaptic signaling & cognition | Morphogenesis and anatomical formations |
| Anion membrane transport | - | - |
| Network **1** *vs* **2**/**3** *(Same as in Fig. 4)* | Network **2** *vs* **1**/**3** | Network **3** *vs* **1**/**2** |

**Supplementary Fig. 13. Whole transcriptome differential gene expression and gene ontology (GO) analysis for each mitochondria-derived brain network.** RNA transcript levels (*in situ* hybridization) were obtained from the Allen Mouse Brain Atlas, and averaged for all brain areas representing the 16 of the 17 punched areas (dorsal and ventral DG not distinguished in the reference dataset) used in the mitochondrial analyses (**Supplemental File 1**). Regional gene expression (n=16) was then averaged for each network (*Network 1* = 6 areas; *Network 2* = 7 areas; *Network 3* = 4 areas). Genes with expression above a Log2-fold difference >1 (double the expression relative to all other areas) were extracted as OVER-expressed, and genes with expression with Log2 fold difference <1 (half the expression relative to all other areas) were extracted as UNDER-expressed. The number of differentially expressed genes is listed in the tables (*right*) and the gene lists available in **Supplemental File 2**. From these gene lists, ShinyGo 0.76.3 was used to identify significantly over- and under-represented biological pathways (FDR<0.05), which were then grouped and analyzed as networks of related pathways, and summarized as categories in the tables above. Only two major categories of pathways were significant for the under-expressed genes among Networks 2 and 3. Source data are provided as a Source Data file.

**Supplemental Table 1.** Study design with the number of animals allocated to each condition (total n=28). Naïve represent untreated control animals for each study group, and days at sacrifice reflect the time elapsed since the onset of experiments.

| Experimental group | Naïve (CORT) | CORT | Naïve (CSDS-matched) | CSDS | Naïve (CSDS Recovered) | CSDS-recovered |
|---|---|---|---|---|---|---|
| Number of animals | 5 | 5 | 3 | 6 | 3 | 6 |
| Day at sacrifice | 63 | 63 | 14 | 14 | 71 | 71 |
