## Supplementary Data 3 for "Brain mitochondrial diversity and network organization predict anxiety-like behavior in mice": Supplementary Data 3 copy.html

Supplementary file 3


Code 

- Show All Code
- Hide All Code

### Supplementary file 3

###### Compiled: March 03, 2023

- Introduction and data
  preparation
- Figure 4b - Principal
  component analysis
- Figure 4c - Hierarchical
  clustering
- Figure 4d - Ranked pathway
  scores
- Figure
  4e - Bivariate plot of Vitamin B2 metabolism & G3P shuttle
- Figure 4f - Ratio G3P/Vit.
  B2
- Figure
  4g - Bivariate plot of Calcium homeostasis & Metabolism
- Figure 4h -
  Ratio Calcium homeostasis/Metabolism

##### Introduction and data preparation

This markdown file contains code and figures shown in Figure 4. Code
chunks can be expanded, and plots were generated interactively where
applicable. This file also contains additional information (tables,
interactive plots) supporting the analysis. We used the Allen mouse
brain atlas dataset and human MitoCarta3.0 to study mitochondrial
signatures in mouse brain tissue. The following datasets were used:

**Allen mouse brain atlas (Harmonizome) dataset (gene
expression from in situ hybridization)**  
https://maayanlab.cloud/Harmonizome/dataset/Allen+Brain+Atlas+Adult+Mouse+Brain+Tissue+Gene+Expression+Profiles

**Mouse and human Mitocarta 3.0 datasets:**  
<https://www.broadinstitute.org/mitocarta/mitocarta30-inventory-mammalian-mitochondrial-proteins-and-pathways>

The Allen dataset was annotated with human (not mouse) entrez IDs,
and was thus mapped to the human MitoCarta dataset. In total, we found
946 mitochondrial genes in the Allen dataset. Among the un-identified
genes were for instance 12 mitochondrial DNA-encoded genes (with the
exception of ND3) and several electron transport chain complex
subunits.

```
## Read the raw data
data_raw <- read.delim("Data/gene_attribute_matrix_cleaned.txt")
## Keep a dataframe with ID and Gene name
gene_symbol  <- data_raw[3:nrow(data_raw),1]
gene_ID  <- data_raw[3:nrow(data_raw),3]
ID_to_symbol <- data.frame(Allen_gene_symbol = gene_symbol, Allen_gene_ID = gene_ID)
## Create dataframe with Structures and Acronyms 
structures <- t(data_raw[1, 4:ncol(data_raw)]) %>%
  as.data.frame() %>%
  rownames_to_column("Structure") %>%
  dplyr::rename(StructureAcronym = "1")
## New dataframe with raw data, structures, acronyms and gene IDs 
data_raw <- data_raw[3:nrow(data_raw),4:ncol(data_raw)] 
rownames(data_raw) <- gene_ID
data_raw  <- data_raw  %>%
  rownames_to_column("ID") %>%
  pivot_longer(cols = colnames(data_raw ), names_to = "Structure", values_to = "exprs") %>%
  mutate(exprs = as.numeric(exprs)) %>%
  full_join(structures, by = "Structure")
## Filter for brain Areas of interest 
Areas_to_keep <- read.csv("Data/Areas_to_keep.csv")
data_raw  <- data_raw  %>%
  filter(StructureAcronym %in% Areas_to_keep$Acronym)
## Filter for mitochondrial genes
gene_to_ID_mitocarta_hm <- readxl::read_xls(here::here("Data", "HumanMitoCarta3_0.xls"), sheet = 2) %>%
  dplyr::select(HumanGeneID,  Symbol) %>%
  unique() %>%
  mutate(HumanGeneID = as.character(HumanGeneID)) 
mitoIDs_hm <- unique(gene_to_ID_mitocarta_hm$HumanGeneID)
data_raw_mito <- data_raw %>%
  dplyr::filter(ID %in% mitoIDs_hm)%>%
  dplyr::mutate(exprs = as.numeric(exprs)) %>%
  pivot_wider(names_from = "ID", values_from = "exprs")
```

**Mitochondrial genes identified in the Allen brain atlas
dataset and included in this analysis:**

```
Mitocarta_in_Allen <- ID_to_symbol %>%
  filter(Allen_gene_ID %in% c(mitoIDs_hm)) %>%
    unique() 
Mitocarta_in_Allen_table <- gene_to_ID_mitocarta_hm %>%
  filter(HumanGeneID %in% Mitocarta_in_Allen$Allen_gene_ID) %>%
  arrange(Symbol)
knitr::kable(Mitocarta_in_Allen_table, caption = "Included") %>% 
  kableExtra::kable_styling(full_width = F) %>%
  kableExtra::scroll_box(width = "500px", height = "400px")
```

Included

| HumanGeneID | Symbol |
| --- | --- |
| 51166 | AADAT |
| 10157 | AASS |
| 18 | ABAT |
| 10350 | ABCA9 |
| 23456 | ABCB10 |
| 10058 | ABCB6 |
| 22 | ABCB7 |
| 11194 | ABCB8 |
| 215 | ABCD1 |
| 225 | ABCD2 |
| 5825 | ABCD3 |
| 55347 | ABHD10 |
| 83451 | ABHD11 |
| 10449 | ACAA2 |
| 31 | ACACA |
| 32 | ACACB |
| 80724 | ACAD10 |
| 84129 | ACAD11 |
| 27034 | ACAD8 |
| 28976 | ACAD9 |
| 33 | ACADL |
| 34 | ACADM |
| 35 | ACADS |
| 36 | ACADSB |
| 37 | ACADVL |
| 38 | ACAT1 |
| 84680 | ACCS |
| 47 | ACLY |
| 50 | ACO2 |
| 55856 | ACOT13 |
| 10965 | ACOT2 |
| 23597 | ACOT9 |
| 51205 | ACP6 |
| 197322 | ACSF3 |
| 2180 | ACSL1 |
| 23305 | ACSL6 |
| 123876 | ACSM2A |
| 6296 | ACSM3 |
| 341392 | ACSM4 |
| 54988 | ACSM5 |
| 84532 | ACSS1 |
| 79611 | ACSS3 |
| 57143 | ADCK1 |
| 90956 | ADCK2 |
| 203054 | ADCK5 |
| 55811 | ADCY10 |
| 137872 | ADHFE1 |
| 246269 | AFG1L |
| 10939 | AFG3L2 |
| 55750 | AGK |
| 79814 | AGMAT |
| 56895 | AGPAT4 |
| 55326 | AGPAT5 |
| 189 | AGXT |
| 64902 | AGXT2 |
| 10768 | AHCYL1 |
| 9131 | AIFM1 |
| 84883 | AIFM2 |
| 150209 | AIFM3 |
| 204 | AK2 |
| 50808 | AK3 |
| 205 | AK4 |
| 8165 | AKAP1 |
| 11216 | AKAP10 |
| 57016 | AKR1B10 |
| 8574 | AKR7A2 |
| 211 | ALAS1 |
| 212 | ALAS2 |
| 5832 | ALDH18A1 |
| 219 | ALDH1B1 |
| 10840 | ALDH1L1 |
| 160428 | ALDH1L2 |
| 217 | ALDH2 |
| 224 | ALDH3A2 |
| 8659 | ALDH4A1 |
| 7915 | ALDH5A1 |
| 4329 | ALDH6A1 |
| 501 | ALDH7A1 |
| 223 | ALDH9A1 |
| 8846 | ALKBH1 |
| 23600 | AMACR |
| 275 | AMT |
| 90806 | ANGEL2 |
| 65990 | ANTKMT |
| 328 | APEX1 |
| 79135 | APOO |
| 139322 | APOOL |
| 381 | ARF5 |
| 384 | ARG2 |
| 402 | ARL2 |
| 83787 | ARMC10 |
| 51309 | ARMCX1 |
| 9823 | ARMCX2 |
| 51566 | ARMCX3 |
| 54470 | ARMCX6 |
| 84896 | ATAD1 |
| 55210 | ATAD3A |
| 91419 | ATP23 |
| 498 | ATP5F1A |
| 509 | ATP5F1C |
| 513 | ATP5F1D |
| 93974 | ATP5IF1 |
| 516 | ATP5MC1 |
| 518 | ATP5MC3 |
| 84833 | ATP5MD |
| 9551 | ATP5MF |
| 100526740 | ATP5MF-PTCD1 |
| 10632 | ATP5MG |
| 9556 | ATP5MPL |
| 515 | ATP5PB |
| 10476 | ATP5PD |
| 522 | ATP5PF |
| 539 | ATP5PO |
| 91647 | ATPAF2 |
| 134145 | ATPSCKMT |
| 549 | AUH |
| 54998 | AURKAIP1 |
| 578 | BAK1 |
| 581 | BAX |
| 27113 | BBC3 |
| 587 | BCAT2 |
| 594 | BCKDHB |
| 10295 | BCKDK |
| 596 | BCL2 |
| 597 | BCL2A1 |
| 598 | BCL2L1 |
| 10017 | BCL2L10 |
| 10018 | BCL2L11 |
| 23786 | BCL2L13 |
| 599 | BCL2L2 |
| 617 | BCS1L |
| 637 | BID |
| 638 | BIK |
| 2647 | BLOC1S1 |
| 664 | BNIP3 |
| 665 | BNIP3L |
| 666 | BOK |
| 51027 | BOLA1 |
| 388962 | BOLA3 |
| 670 | BPHL |
| 91574 | C12orf65 |
| 84419 | C15orf48 |
| 145853 | C15orf61 |
| 283951 | C16orf91 |
| 708 | C1QBP |
| 205327 | C2orf69 |
| 285315 | C3orf33 |
| 401207 | C5orf63 |
| 221545 | C6orf136 |
| 414919 | C8orf82 |
| 763 | CA5A |
| 11238 | CA5B |
| 836 | CASP3 |
| 841 | CASP8 |
| 842 | CASP9 |
| 847 | CAT |
| 874 | CBR3 |
| 84869 | CBR4 |
| 133957 | CCDC127 |
| 79714 | CCDC51 |
| 131076 | CCDC58 |
| 60492 | CCDC90B |
| 51654 | CDK5RAP1 |
| 84902 | CEP89 |
| 118487 | CHCHD1 |
| 400916 | CHCHD10 |
| 51142 | CHCHD2 |
| 54927 | CHCHD3 |
| 131474 | CHCHD4 |
| 84269 | CHCHD5 |
| 84303 | CHCHD6 |
| 79145 | CHCHD7 |
| 55349 | CHDH |
| 56994 | CHPT1 |
| 55847 | CISD1 |
| 284106 | CISD3 |
| 1160 | CKMT2 |
| 81570 | CLPB |
| 8192 | CLPP |
| 10845 | CLPX |
| 171425 | CLYBL |
| 152100 | CMC1 |
| 56942 | CMC2 |
| 129607 | CMPK2 |
| 28958 | COA3 |
| 51287 | COA4 |
| 493753 | COA5 |
| 84334 | COA8 |
| 80347 | COASY |
| 1312 | COMT |
| 118881 | COMTD1 |
| 93058 | COQ10A |
| 80219 | COQ10B |
| 51805 | COQ3 |
| 51117 | COQ4 |
| 84274 | COQ5 |
| 51004 | COQ6 |
| 10229 | COQ7 |
| 56997 | COQ8A |
| 79934 | COQ8B |
| 57017 | COQ9 |
| 1352 | COX10 |
| 1353 | COX11 |
| 84987 | COX14 |
| 1355 | COX15 |
| 51241 | COX16 |
| 10063 | COX17 |
| 285521 | COX18 |
| 90639 | COX19 |
| 116228 | COX20 |
| 1327 | COX4I1 |
| 84701 | COX4I2 |
| 9377 | COX5A |
| 1337 | COX6A1 |
| 1340 | COX6B1 |
| 125965 | COX6B2 |
| 1346 | COX7A1 |
| 1347 | COX7A2 |
| 9167 | COX7A2L |
| 1349 | COX7B |
| 1371 | CPOX |
| 1373 | CPS1 |
| 1374 | CPT1A |
| 1375 | CPT1B |
| 126129 | CPT1C |
| 1384 | CRAT |
| 54675 | CRLS1 |
| 54677 | CROT |
| 1407 | CRY1 |
| 1429 | CRYZ |
| 1431 | CS |
| 80777 | CYB5B |
| 1537 | CYC1 |
| 54205 | CYCS |
| 1583 | CYP11A1 |
| 1591 | CYP24A1 |
| 1593 | CYP27A1 |
| 1594 | CYP27B1 |
| 728294 | D2HGDH |
| 7818 | DAP3 |
| 55157 | DARS2 |
| 1622 | DBI |
| 1629 | DBT |
| 79877 | DCAKD |
| 51181 | DCXR |
| 55794 | DDX28 |
| 1666 | DECR1 |
| 9812 | DELE1 |
| 80017 | DGLUCY |
| 1716 | DGUOK |
| 115817 | DHRS1 |
| 10901 | DHRS4 |
| 25979 | DHRS7B |
| 55526 | DHTKD1 |
| 22907 | DHX30 |
| 56616 | DIABLO |
| 1737 | DLAT |
| 1743 | DLST |
| 90871 | DMAC1 |
| 55101 | DMAC2 |
| 27109 | DMAC2L |
| 29958 | DMGDH |
| 1763 | DNA2 |
| 9093 | DNAJA3 |
| 55735 | DNAJC11 |
| 29103 | DNAJC15 |
| 131118 | DNAJC19 |
| 54943 | DNAJC28 |
| 3338 | DNAJC4 |
| 728489 | DNLZ |
| 10059 | DNM1L |
| 1841 | DTYMK |
| 54920 | DUS2 |
| 1854 | DUT |
| 124454 | EARS2 |
| 1891 | ECH1 |
| 55862 | ECHDC1 |
| 55268 | ECHDC2 |
| 1892 | ECHS1 |
| 1632 | ECI1 |
| 10455 | ECI2 |
| 51295 | ECSIT |
| 80303 | EFHD1 |
| 1962 | EHHADH |
| 60528 | ELAC2 |
| 2021 | ENDOG |
| 2053 | EPHX2 |
| 26284 | ERAL1 |
| 2108 | ETFA |
| 2109 | ETFB |
| 254013 | ETFBKMT |
| 2110 | ETFDH |
| 144363 | ETFRF1 |
| 23474 | ETHE1 |
| 55218 | EXD2 |
| 9941 | EXOG |
| 2168 | FABP1 |
| 81889 | FAHD1 |
| 84908 | FAM136A |
| 26355 | FAM162A |
| 222234 | FAM185A |
| 125228 | FAM210A |
| 116151 | FAM210B |
| 10667 | FARS2 |
| 2194 | FASN |
| 10922 | FASTK |
| 79675 | FASTKD1 |
| 22868 | FASTKD2 |
| 79072 | FASTKD3 |
| 60493 | FASTKD5 |
| 26235 | FBXL4 |
| 2224 | FDPS |
| 2230 | FDX1 |
| 2232 | FDXR |
| 2235 | FECH |
| 2271 | FH |
| 2272 | FHIT |
| 51024 | FIS1 |
| 60681 | FKBP10 |
| 23770 | FKBP8 |
| 80308 | FLAD1 |
| 154791 | FMC1 |
| 2356 | FPGS |
| 2495 | FTH1 |
| 94033 | FTMT |
| 139341 | FUNDC1 |
| 65991 | FUNDC2 |
| 2395 | FXN |
| 90480 | GADD45GIP1 |
| 2617 | GARS1 |
| 5188 | GATB |
| 283459 | GATC |
| 2628 | GATM |
| 23464 | GCAT |
| 2639 | GCDH |
| 2671 | GFER |
| 84340 | GFM2 |
| 27069 | GHITM |
| 2731 | GLDC |
| 51031 | GLOD4 |
| 51022 | GLRX2 |
| 2744 | GLS |
| 27165 | GLS2 |
| 2746 | GLUD1 |
| 10249 | GLYAT |
| 132158 | GLYCTK |
| 64083 | GOLPH3 |
| 2806 | GOT2 |
| 57678 | GPAM |
| 150763 | GPAT2 |
| 2820 | GPD2 |
| 84706 | GPT2 |
| 2876 | GPX1 |
| 2879 | GPX4 |
| 9380 | GRHPR |
| 134266 | GRPEL2 |
| 2926 | GRSF1 |
| 2936 | GSR |
| 373156 | GSTK1 |
| 2954 | GSTZ1 |
| 85865 | GTPBP10 |
| 84705 | GTPBP3 |
| 60558 | GUF1 |
| 2987 | GUK1 |
| 3033 | HADH |
| 3030 | HADHA |
| 3032 | HADHB |
| 3029 | HAGH |
| 51179 | HAO2 |
| 23438 | HARS2 |
| 3052 | HCCS |
| 81932 | HDHD3 |
| 50865 | HEBP1 |
| 51409 | HEMK1 |
| 26275 | HIBCH |
| 25994 | HIGD1A |
| 192286 | HIGD2A |
| 84681 | HINT2 |
| 135114 | HINT3 |
| 3155 | HMGCL |
| 3158 | HMGCS2 |
| 112817 | HOGA1 |
| 84842 | HPDL |
| 150274 | HSCB |
| 3028 | HSD17B10 |
| 3295 | HSD17B4 |
| 83693 | HSDL1 |
| 3313 | HSPA9 |
| 3329 | HSPD1 |
| 3336 | HSPE1 |
| 10553 | HTATIP2 |
| 27429 | HTRA2 |
| 55699 | IARS2 |
| 200205 | IBA57 |
| 3416 | IDE |
| 3418 | IDH2 |
| 3419 | IDH3A |
| 3420 | IDH3B |
| 3421 | IDH3G |
| 3422 | IDI1 |
| 83943 | IMMP2L |
| 81689 | ISCA1 |
| 122961 | ISCA2 |
| 23479 | ISCU |
| 3712 | IVD |
| 3735 | KARS1 |
| 8564 | KMO |
| 56267 | KYAT3 |
| 79944 | L2HGDH |
| 114294 | LACTB |
| 51110 | LACTB2 |
| 51056 | LAP3 |
| 23395 | LARS2 |
| 197257 | LDHD |
| 3954 | LETM1 |
| 137994 | LETM2 |
| 25875 | LETMD1 |
| 11019 | LIAS |
| 3980 | LIG3 |
| 51601 | LIPT1 |
| 387787 | LIPT2 |
| 9361 | LONP1 |
| 10128 | LRPPRC |
| 10434 | LYPLA1 |
| 127018 | LYPLAL1 |
| 57149 | LYRM1 |
| 57226 | LYRM2 |
| 57128 | LYRM4 |
| 201229 | LYRM9 |
| 28992 | MACROD1 |
| 79568 | MAIP1 |
| 115416 | MALSU1 |
| 4128 | MAOA |
| 4129 | MAOB |
| 54708 | MARCHF5 |
| 92935 | MARS2 |
| 57506 | MAVS |
| 27349 | MCAT |
| 56922 | MCCC1 |
| 64087 | MCCC2 |
| 84693 | MCEE |
| 4170 | MCL1 |
| 84331 | MCRIP2 |
| 90550 | MCU |
| 55013 | MCUB |
| 63933 | MCUR1 |
| 4200 | ME2 |
| 10873 | ME3 |
| 51102 | MECR |
| 254042 | METAP1D |
| 196074 | METTL15 |
| 64863 | METTL4 |
| 29081 | METTL5 |
| 79828 | METTL8 |
| 9927 | MFN2 |
| 84709 | MGARP |
| 92667 | MGME1 |
| 4259 | MGST3 |
| 440574 | MICOS10 |
| 125988 | MICOS13 |
| 10367 | MICU1 |
| 221154 | MICU2 |
| 286097 | MICU3 |
| 54471 | MIEF1 |
| 125170 | MIEF2 |
| 374986 | MIGA1 |
| 23417 | MLYCD |
| 166785 | MMAA |
| 27249 | MMADHC |
| 4594 | MMUT |
| 51660 | MPC1 |
| 25874 | MPC2 |
| 4357 | MPST |
| 4358 | MPV17 |
| 255027 | MPV17L |
| 84769 | MPV17L2 |
| 79922 | MRM1 |
| 29960 | MRM2 |
| 55178 | MRM3 |
| 65008 | MRPL1 |
| 124995 | MRPL10 |
| 65003 | MRPL11 |
| 6182 | MRPL12 |
| 28998 | MRPL13 |
| 54948 | MRPL16 |
| 63875 | MRPL17 |
| 29074 | MRPL18 |
| 9801 | MRPL19 |
| 55052 | MRPL20 |
| 219927 | MRPL21 |
| 29093 | MRPL22 |
| 79590 | MRPL24 |
| 51264 | MRPL27 |
| 11222 | MRPL3 |
| 51263 | MRPL30 |
| 64983 | MRPL32 |
| 9553 | MRPL33 |
| 64981 | MRPL34 |
| 51318 | MRPL35 |
| 64979 | MRPL36 |
| 51253 | MRPL37 |
| 64978 | MRPL38 |
| 54148 | MRPL39 |
| 51073 | MRPL4 |
| 64976 | MRPL40 |
| 64975 | MRPL41 |
| 84545 | MRPL43 |
| 65080 | MRPL44 |
| 84311 | MRPL45 |
| 26589 | MRPL46 |
| 57129 | MRPL47 |
| 51642 | MRPL48 |
| 740 | MRPL49 |
| 54534 | MRPL50 |
| 51258 | MRPL51 |
| 122704 | MRPL52 |
| 116540 | MRPL53 |
| 116541 | MRPL54 |
| 128308 | MRPL55 |
| 78988 | MRPL57 |
| 3396 | MRPL58 |
| 65005 | MRPL9 |
| 55173 | MRPS10 |
| 64963 | MRPS11 |
| 6183 | MRPS12 |
| 64960 | MRPS15 |
| 51021 | MRPS16 |
| 51373 | MRPS17 |
| 55168 | MRPS18A |
| 28973 | MRPS18B |
| 51023 | MRPS18C |
| 51116 | MRPS2 |
| 54460 | MRPS21 |
| 56945 | MRPS22 |
| 51649 | MRPS23 |
| 64951 | MRPS24 |
| 64432 | MRPS25 |
| 28957 | MRPS28 |
| 10240 | MRPS31 |
| 51650 | MRPS33 |
| 65993 | MRPS34 |
| 60488 | MRPS35 |
| 92259 | MRPS36 |
| 64969 | MRPS5 |
| 64968 | MRPS6 |
| 51081 | MRPS7 |
| 64965 | MRPS9 |
| 92399 | MRRF |
| 57380 | MRS2 |
| 22921 | MSRB2 |
| 253827 | MSRB3 |
| 4537 | MT-ND3 |
| 64757 | MTARC1 |
| 54996 | MTARC2 |
| 23787 | MTCH1 |
| 23788 | MTCH2 |
| 7978 | MTERF1 |
| 80298 | MTERF2 |
| 51001 | MTERF3 |
| 130916 | MTERF4 |
| 51537 | MTFP1 |
| 56181 | MTFR1L |
| 26164 | MTG2 |
| 25902 | MTHFD1L |
| 10797 | MTHFD2 |
| 441024 | MTHFD2L |
| 10588 | MTHFS |
| 4528 | MTIF2 |
| 219402 | MTIF3 |
| 25821 | MTO1 |
| 55149 | MTPAP |
| 51250 | MTRES1 |
| 54516 | MTRF1L |
| 4580 | MTX1 |
| 10651 | MTX2 |
| 79594 | MUL1 |
| 4595 | MUTYH |
| 60314 | MYG1 |
| 133686 | NADK2 |
| 162417 | NAGS |
| 79731 | NARS2 |
| 339983 | NAT8L |
| 128240 | NAXE |
| 4077 | NBR1 |
| 4694 | NDUFA1 |
| 4705 | NDUFA10 |
| 55967 | NDUFA12 |
| 4695 | NDUFA2 |
| 4696 | NDUFA3 |
| 4697 | NDUFA4 |
| 4698 | NDUFA5 |
| 4700 | NDUFA6 |
| 4701 | NDUFA7 |
| 4702 | NDUFA8 |
| 4704 | NDUFA9 |
| 4706 | NDUFAB1 |
| 51103 | NDUFAF1 |
| 91942 | NDUFAF2 |
| 25915 | NDUFAF3 |
| 29078 | NDUFAF4 |
| 79133 | NDUFAF5 |
| 55471 | NDUFAF7 |
| 284184 | NDUFAF8 |
| 4716 | NDUFB10 |
| 4708 | NDUFB2 |
| 4709 | NDUFB3 |
| 4710 | NDUFB4 |
| 4711 | NDUFB5 |
| 4712 | NDUFB6 |
| 4713 | NDUFB7 |
| 4714 | NDUFB8 |
| 4715 | NDUFB9 |
| 4717 | NDUFC1 |
| 4718 | NDUFC2 |
| 4719 | NDUFS1 |
| 4722 | NDUFS3 |
| 4724 | NDUFS4 |
| 4726 | NDUFS6 |
| 4728 | NDUFS8 |
| 4723 | NDUFV1 |
| 4729 | NDUFV2 |
| 4731 | NDUFV3 |
| 129807 | NEU4 |
| 9054 | NFS1 |
| 27247 | NFU1 |
| 51335 | NGRN |
| 60491 | NIF3L1 |
| 8508 | NIPSNAP1 |
| 2631 | NIPSNAP2 |
| 25934 | NIPSNAP3A |
| 4817 | NIT1 |
| 56954 | NIT2 |
| 57486 | NLN |
| 79671 | NLRX1 |
| 4832 | NME3 |
| 4833 | NME4 |
| 10201 | NME6 |
| 25819 | NOCT |
| 54888 | NSUN2 |
| 63899 | NSUN3 |
| 387338 | NSUN4 |
| 64943 | NT5DC2 |
| 51559 | NT5DC3 |
| 56953 | NT5M |
| 80224 | NUBPL |
| 25961 | NUDT13 |
| 390916 | NUDT19 |
| 318 | NUDT2 |
| 11164 | NUDT5 |
| 53343 | NUDT9 |
| 4942 | OAT |
| 54940 | OCIAD1 |
| 132299 | OCIAD2 |
| 4967 | OGDH |
| 55753 | OGDHL |
| 4968 | OGG1 |
| 115209 | OMA1 |
| 4976 | OPA1 |
| 80207 | OPA3 |
| 114876 | OSBPL1A |
| 64172 | OSGEPL1 |
| 5009 | OTC |
| 5018 | OXA1L |
| 5019 | OXCT1 |
| 339229 | OXLD1 |
| 92106 | OXNAD1 |
| 55074 | OXR1 |
| 54995 | OXSM |
| 140886 | PABPC5 |
| 10606 | PAICS |
| 51025 | PAM16 |
| 5091 | PC |
| 84105 | PCBD2 |
| 5095 | PCCA |
| 5096 | PCCB |
| 5106 | PCK2 |
| 201626 | PDE12 |
| 5160 | PDHA1 |
| 5162 | PDHB |
| 8050 | PDHX |
| 5163 | PDK1 |
| 5165 | PDK3 |
| 5166 | PDK4 |
| 54704 | PDP1 |
| 57546 | PDP2 |
| 23590 | PDSS1 |
| 57107 | PDSS2 |
| 8799 | PEX11B |
| 192111 | PGAM5 |
| 9489 | PGS1 |
| 5245 | PHB |
| 11331 | PHB2 |
| 5264 | PHYH |
| 9463 | PICK1 |
| 80119 | PIF1 |
| 23761 | PISD |
| 10531 | PITRM1 |
| 55848 | PLGRKT |
| 11212 | PLPBP |
| 57048 | PLSCR3 |
| 23203 | PMPCA |
| 25953 | PNKD |
| 55163 | PNPO |
| 87178 | PNPT1 |
| 5423 | POLB |
| 26073 | POLDIP2 |
| 5428 | POLG |
| 11232 | POLG2 |
| 10721 | POLQ |
| 5442 | POLRMT |
| 27068 | PPA2 |
| 10105 | PPIF |
| 152926 | PPM1K |
| 5498 | PPOX |
| 160760 | PPTC7 |
| 7001 | PRDX2 |
| 10549 | PRDX4 |
| 25824 | PRDX5 |
| 9588 | PRDX6 |
| 27166 | PRELID1 |
| 153768 | PRELID2 |
| 10650 | PRELID3A |
| 51012 | PRELID3B |
| 9581 | PREPL |
| 201973 | PRIMPOL |
| 5071 | PRKN |
| 58510 | PRODH2 |
| 9692 | PRORP |
| 167681 | PRSS35 |
| 84293 | PRXL2A |
| 79810 | PTCD2 |
| 55037 | PTCD3 |
| 114971 | PTPMT1 |
| 138428 | PTRH1 |
| 51651 | PTRH2 |
| 126789 | PUSL1 |
| 5827 | PXMP2 |
| 11264 | PXMP4 |
| 5831 | PYCR1 |
| 29920 | PYCR2 |
| 5860 | QDPR |
| 55278 | QRSL1 |
| 81890 | QTRT1 |
| 53917 | RAB24 |
| 55969 | RAB5IF |
| 57038 | RARS2 |
| 79863 | RBFA |
| 81554 | RCC1L |
| 57665 | RDH14 |
| 9401 | RECQL4 |
| 25996 | REXO2 |
| 55312 | RFK |
| 55288 | RHOT1 |
| 89941 | RHOT2 |
| 10247 | RIDA |
| 51115 | RMDN1 |
| 55177 | RMDN3 |
| 55005 | RMND1 |
| 246243 | RNASEH1 |
| 140823 | ROMO1 |
| 84881 | RPUSD4 |
| 55316 | RSAD1 |
| 84816 | RTN4IP1 |
| 25813 | SAMM50 |
| 1757 | SARDH |
| 54938 | SARS2 |
| 6341 | SCO1 |
| 6342 | SCP2 |
| 6389 | SDHA |
| 54949 | SDHAF2 |
| 57001 | SDHAF3 |
| 6390 | SDHB |
| 6391 | SDHC |
| 6392 | SDHD |
| 56948 | SDR39U1 |
| 113675 | SDSL |
| 83642 | SELENOO |
| 5414 | SEPTIN4 |
| 84947 | SERAC1 |
| 94081 | SFXN1 |
| 118980 | SFXN2 |
| 81855 | SFXN3 |
| 119559 | SFXN4 |
| 94097 | SFXN5 |
| 6472 | SHMT2 |
| 23410 | SIRT3 |
| 23409 | SIRT4 |
| 23408 | SIRT5 |
| 1468 | SLC25A10 |
| 8402 | SLC25A11 |
| 8604 | SLC25A12 |
| 10165 | SLC25A13 |
| 9016 | SLC25A14 |
| 8034 | SLC25A16 |
| 83733 | SLC25A18 |
| 60386 | SLC25A19 |
| 788 | SLC25A20 |
| 89874 | SLC25A21 |
| 79751 | SLC25A22 |
| 79085 | SLC25A23 |
| 29957 | SLC25A24 |
| 114789 | SLC25A25 |
| 115286 | SLC25A26 |
| 9481 | SLC25A27 |
| 123096 | SLC25A29 |
| 5250 | SLC25A3 |
| 253512 | SLC25A30 |
| 83447 | SLC25A31 |
| 81034 | SLC25A32 |
| 84275 | SLC25A33 |
| 284723 | SLC25A34 |
| 399512 | SLC25A35 |
| 55186 | SLC25A36 |
| 51312 | SLC25A37 |
| 54977 | SLC25A38 |
| 51629 | SLC25A39 |
| 55972 | SLC25A40 |
| 284427 | SLC25A41 |
| 284439 | SLC25A42 |
| 203427 | SLC25A43 |
| 9673 | SLC25A44 |
| 283130 | SLC25A45 |
| 91137 | SLC25A46 |
| 292 | SLC25A5 |
| 92014 | SLC25A51 |
| 401612 | SLC25A53 |
| 10463 | SLC30A9 |
| 80024 | SLC8B1 |
| 81892 | SLIRP |
| 389203 | SMIM20 |
| 57150 | SMIM8 |
| 9342 | SNAP29 |
| 27044 | SND1 |
| 6647 | SOD1 |
| 219938 | SPATA19 |
| 64847 | SPATA20 |
| 6687 | SPG7 |
| 56848 | SPHK2 |
| 80309 | SPHKAP |
| 56907 | SPIRE1 |
| 6697 | SPR |
| 283377 | SPRYD4 |
| 9517 | SPTLC2 |
| 58472 | SQOR |
| 6742 | SSBP1 |
| 6770 | STAR |
| 56910 | STARD7 |
| 2040 | STOM |
| 30968 | STOML2 |
| 55014 | STX17 |
| 51657 | STYXL1 |
| 8803 | SUCLA2 |
| 8802 | SUCLG1 |
| 8801 | SUCLG2 |
| 79783 | SUGCT |
| 6821 | SUOX |
| 6832 | SUPV3L1 |
| 6834 | SURF1 |
| 55333 | SYNJ2BP |
| 132001 | TAMM41 |
| 80222 | TARS2 |
| 6901 | TAZ |
| 9238 | TBRG4 |
| 285343 | TCAIM |
| 79736 | TEFM |
| 7019 | TFAM |
| 51106 | TFB1M |
| 117145 | THEM4 |
| 284486 | THEM5 |
| 54974 | THG1L |
| 79896 | THNSL1 |
| 26519 | TIMM10 |
| 26515 | TIMM10B |
| 26517 | TIMM13 |
| 10440 | TIMM17A |
| 10245 | TIMM17B |
| 29090 | TIMM21 |
| 29928 | TIMM22 |
| 100287932 | TIMM23 |
| 90580 | TIMM29 |
| 10469 | TIMM44 |
| 92609 | TIMM50 |
| 1678 | TIMM8A |
| 26521 | TIMM8B |
| 26520 | TIMM9 |
| 51300 | TIMMDC1 |
| 8834 | TMEM11 |
| 84233 | TMEM126A |
| 55863 | TMEM126B |
| 55260 | TMEM143 |
| 51522 | TMEM14C |
| 80775 | TMEM177 |
| 25880 | TMEM186 |
| 374882 | TMEM205 |
| 157378 | TMEM65 |
| 55217 | TMLHE |
| 9804 | TOMM20 |
| 56993 | TOMM22 |
| 10953 | TOMM34 |
| 10452 | TOMM40 |
| 84134 | TOMM40L |
| 100188893 | TOMM6 |
| 9868 | TOMM70 |
| 116447 | TOP1MT |
| 7156 | TOP3A |
| 10131 | TRAP1 |
| 51499 | TRIAP1 |
| 54802 | TRIT1 |
| 55621 | TRMT1 |
| 54931 | TRMT10C |
| 79979 | TRMT2B |
| 57570 | TRMT5 |
| 55687 | TRMU |
| 51095 | TRNT1 |
| 26995 | TRUB2 |
| 10102 | TSFM |
| 706 | TSPO |
| 7263 | TST |
| 100131187 | TSTD1 |
| 54902 | TTC19 |
| 7284 | TUFM |
| 56652 | TWNK |
| 25828 | TXN2 |
| 7296 | TXNRD1 |
| 10587 | TXNRD2 |
| 7350 | UCP1 |
| 7351 | UCP2 |
| 7352 | UCP3 |
| 7374 | UNG |
| 84300 | UQCC2 |
| 29796 | UQCR10 |
| 10975 | UQCR11 |
| 7384 | UQCRC1 |
| 7385 | UQCRC2 |
| 7386 | UQCRFS1 |
| 27089 | UQCRQ |
| 84749 | USP30 |
| 57176 | VARS2 |
| 7416 | VDAC1 |
| 7419 | VDAC3 |
| 23078 | VWA8 |
| 10352 | WARS2 |
| 51067 | YARS2 |
| 54059 | YBEY |
| 10730 | YME1L1 |
| 79693 | YRDC |

**Mitochondrial genes not identified in the Allen brain atlas
dataset and not included in this analysis:**

```
Mitocarta_not_in_Allen <- gene_to_ID_mitocarta_hm %>%
  filter(!HumanGeneID %in% Mitocarta_in_Allen$Allen_gene_ID) %>%
  arrange(Symbol)
knitr::kable(Mitocarta_not_in_Allen, caption = "Not included") %>% 
  kableExtra::kable_styling(full_width = F) %>%
  kableExtra::scroll_box(width = "500px", height = "400px")
```

Not included

| HumanGeneID | Symbol |
| --- | --- |
| 57505 | AARS2 |
| 30 | ACAA1 |
| 730249 | ACOD1 |
| 26027 | ACOT11 |
| 11332 | ACOT7 |
| 80221 | ACSF2 |
| 116285 | ACSM1 |
| 348158 | ACSM2B |
| 84266 | ALKBH7 |
| 83858 | ATAD3B |
| 506 | ATP5F1B |
| 514 | ATP5F1E |
| 517 | ATP5MC2 |
| 521 | ATP5ME |
| 64756 | ATPAF1 |
| 572 | BAD |
| 593 | BCKDHA |
| 83875 | BCO2 |
| 622 | BDH1 |
| 79587 | CARS2 |
| 548596 | CKMT1A |
| 1159 | CKMT1B |
| 100272147 | CMC4 |
| 55744 | COA1 |
| 388753 | COA6 |
| 65260 | COA7 |
| 27235 | COQ2 |
| 1329 | COX5B |
| 1339 | COX6A2 |
| 1345 | COX6C |
| 170712 | COX7B2 |
| 1350 | COX7C |
| 1351 | COX8A |
| 341947 | COX8C |
| 1376 | CPT2 |
| 751071 | CSKMT |
| 1727 | CYB5R3 |
| 1584 | CYP11B1 |
| 1585 | CYP11B2 |
| 1723 | DHODH |
| 10202 | DHRS2 |
| 1738 | DLD |
| 1760 | DMPK |
| 84277 | DNAJC30 |
| 79746 | ECHDC3 |
| 51011 | FAHD2A |
| 112812 | FDX2 |
| 55572 | FOXRED1 |
| 8209 | GATD3A |
| 2653 | GCSH |
| 54332 | GDAP1 |
| 85476 | GFM1 |
| 51218 | GLRX5 |
| 2747 | GLUD2 |
| 80273 | GRPEL1 |
| 8225 | GTPBP6 |
| 27440 | HDHD5 |
| 11112 | HIBADH |
| 3094 | HINT1 |
| 7923 | HSD17B8 |
| 84263 | HSDL2 |
| 109703458 | HTD2 |
| 3429 | IFI27 |
| 196294 | IMMP1L |
| 10989 | IMMT |
| 79763 | ISOC2 |
| 92483 | LDHAL6B |
| 3945 | LDHB |
| 90624 | LYRM7 |
| 401250 | MCCD1 |
| 4191 | MDH2 |
| 64745 | METTL17 |
| 56947 | MFF |
| 55669 | MFN1 |
| 4257 | MGST1 |
| 84895 | MIGA2 |
| 4285 | MIPEP |
| 326625 | MMAB |
| 4337 | MOCS1 |
| 347411 | MPC1L |
| 64928 | MRPL14 |
| 29088 | MRPL15 |
| 51069 | MRPL2 |
| 6150 | MRPL23 |
| 10573 | MRPL28 |
| 28977 | MRPL42 |
| 63931 | MRPS14 |
| 64949 | MRPS26 |
| 23107 | MRPS27 |
| 10884 | MRPS30 |
| 4482 | MSRA |
| 4508 | MT-ATP6 |
| 4509 | MT-ATP8 |
| 4512 | MT-CO1 |
| 4513 | MT-CO2 |
| 4514 | MT-CO3 |
| 4519 | MT-CYB |
| 4535 | MT-ND1 |
| 4536 | MT-ND2 |
| 4538 | MT-ND4 |
| 4539 | MT-ND4L |
| 4540 | MT-ND5 |
| 4541 | MT-ND6 |
| 123263 | MTFMT |
| 9650 | MTFR1 |
| 113115 | MTFR2 |
| 92170 | MTG1 |
| 9617 | MTRF1 |
| 345778 | MTX3 |
| 80179 | MYO19 |
| 55739 | NAXD |
| 126328 | NDUFA11 |
| 51079 | NDUFA13 |
| 137682 | NDUFAF6 |
| 4707 | NDUFB1 |
| 54539 | NDUFB11 |
| 4720 | NDUFS2 |
| 4725 | NDUFS5 |
| 374291 | NDUFS7 |
| 55335 | NIPSNAP3B |
| 349565 | NMNAT3 |
| 23530 | NNT |
| 84273 | NOA1 |
| 4898 | NRDC |
| 4913 | NTHL1 |
| 11162 | NUDT6 |
| 254552 | NUDT8 |
| 64064 | OXCT2 |
| 80025 | PANK2 |
| 11315 | PARK7 |
| 55486 | PARL |
| 25973 | PARS2 |
| 5138 | PDE2A |
| 64146 | PDF |
| 5161 | PDHA2 |
| 5164 | PDK2 |
| 55066 | PDPR |
| 100131801 | PET100 |
| 100303755 | PET117 |
| 101928527 | PIGBOS1 |
| 65018 | PINK1 |
| 201164 | PLD6 |
| 5366 | PMAIP1 |
| 9512 | PMPCB |
| 50640 | PNPLA8 |
| 10935 | PRDX3 |
| 5566 | PRKACA |
| 5625 | PRODH |
| 26024 | PTCD1 |
| 80142 | PTGES2 |
| 80324 | PUS1 |
| 100996939 | PYURF |
| 112724 | RDH13 |
| NA | RP11\_469A15.2 |
| 22934 | RPIA |
| 285367 | RPUSD3 |
| 79680 | RTL10 |
| 9997 | SCO2 |
| 644096 | SDHAF1 |
| 135154 | SDHAF4 |
| 253190 | SERHL2 |
| 133383 | SETD9 |
| 6576 | SLC25A1 |
| 10166 | SLC25A15 |
| 81894 | SLC25A28 |
| 291 | SLC25A4 |
| 283600 | SLC25A47 |
| 153328 | SLC25A48 |
| 147407 | SLC25A52 |
| 293 | SLC25A6 |
| 91689 | SMDT1 |
| 6648 | SOD2 |
| 51204 | TACO1 |
| 11022 | TDRKH |
| 64216 | TFB2M |
| 7084 | TK2 |
| 54968 | TMEM70 |
| 387990 | TOMM20L |
| 401505 | TOMM5 |
| 54543 | TOMM7 |
| 55006 | TRMT61B |
| 100130890 | TSTD3 |
| 55245 | UQCC1 |
| 790955 | UQCC3 |
| 7381 | UQCRB |
| 7388 | UQCRH |
| 7417 | VDAC2 |
| 55187 | VPS13D |
| 63929 | XPNPEP3 |
| 284273 | ZADH2 |

```
rm(list = setdiff(ls(), c("mitoIDs_hm", "data_raw", "data_mito_raw", "color_groups")))
```

The Allen dataset contains >2000 brain (sub-)areas. To match the
Allen dataset with our mouse dataset, we calculated an average
expression value for each greater area. For instance, the substantia
nigra expression value was averaged across the pars compacta and pars
reticulata. We applied this for all 16 main areas (mOFC, VTA, SN, DG,
PAG, Cereb, VN, mPFC, CPu, NAc, M1, Hypoth, Thal, Amyg, CA3, and V1).
Compared to our dataset with 17 main areas, the Allen dataset did not
divide the dentate gyrus into ventral and dorsal, hence the resulting 16
areas.

The following code describes how the main areas were averaged, and
the table shows the 16 main-areas and the anatomical sub-areas they are
composed of.

```
mainAreas <- data_raw %>%
  dplyr::mutate(Brain_Area = case_when(
    StructureAcronym %in% c("ORBm6a","ORBm2","ORBm1","ORBm2/3","ORBm","ORBm5", "ORB") 
    ~ "mOFC",
    StructureAcronym %in% c("VTA") 
    ~ "VTA",
    StructureAcronym %in% c("SNr", "SNc") 
    ~ "SN",
    StructureAcronym %in% c("DG-mo","DG","DGMol","DG-sg","DGGran","DG-po","DGs","DGi","DGHil") 
    ~ "DG",
    StructureAcronym %in% c("PcPL-PAG", "JcPL-PAG", "PcPV-PAG", "JcPV-PAG", "CoPV-PAG", "m1AD-PAG", 
                            "PIsD-PAG", "p1Lim-PAG", "TGDL-PAG", "TGL-PAG", "SCL-PAG", "SCDL-PAG", 
                            "m1Lim-PAG","ICDL-PAG", "PIsDL-PAG", "PB-PAG", "PIsL-PAG", "isLim-PAG",
                            "m1B-PAG", "p1B-PAG", "Ist-PAG", "PAG") 
    ~ "PAG",
    StructureAcronym %in% c("ANcr1", "ANcr1gr", "ANcr1mo", "CB") 
    ~ "Cereb",
    StructureAcronym %in% c("MV", "LAV", "SPIV", "SUV", "VNC") 
    ~ "VN",
    StructureAcronym %in% c("ILA6b", "ILA", "ILA2/3", "ILA5", "ILA2", "ILA1", 
                            "ILA6a", "PL6b", "PL6a", 
                            "PL1", "PL2/3", "PL", "PL2", "PL5", "ACAd5", "ACAd2/3", 
                            "ACAd","ACA", "ACAd1", "ACAv2/3", "ACAv", "ACAv1", "ACAv5", 
                            "ACAv6a", "ILA",
                            "ACAd6a", "ACAv6b", "ACAd6b", "CCx") 
    ~ "mPFC",
    StructureAcronym %in% c("STRd", "Cau","CP") 
    ~ "CPu",
    StructureAcronym %in% c("AcbSh", "AcbCo", "VStr") 
    ~ "NAc",
    StructureAcronym %in% c("MOp1", "MOp2/3", "MOp5", "MOp6b", "MOp6a", "MOp") 
    ~ "M1",
    StructureAcronym %in% c("PVHIp", "PVHd", "PVHpm", "PVHpml", "PVHm", "PVHmm", "PVHmpd", "PVHmdp", 
                            "PVHp", "PVHap", "PVH", "PVHlp") 
    ~ "Hypoth",
    StructureAcronym %in% c("CL", "CM", "MDc", "MD", "MED", "MDI", "ILM", "MDm", "PVT", "TH", "MDl")
    ~ "Thal",
    StructureAcronym %in% c("BLA", "BLAa", "BLP", "BLAp", "BLA") 
    ~ "Amyg",
    StructureAcronym %in% c("BMAp", "BMP", "BLAv", "BMAa", "BMA")
    ~ "Amyg",
    StructureAcronym %in% c("CA3sp", "CA3sr", "CA3slu", "CA3so", "CA3slm", "CA3")
    ~ "CA3",
    StructureAcronym %in% c("VISp4", "VISp1", "VISp2/3", "VISp6a", "VISp6b", "VISp") 
    ~ "V1"
  )) %>%
  dplyr::mutate(Group = case_when(
    Brain_Area == "mOFC" ~ "Cortico-striatal",
    Brain_Area == "VTA" ~ "Salience/Spatial navigation",
    Brain_Area == "DG" ~ "Salience/Spatial navigation",
    Brain_Area == "PAG" ~ "Threat response",
    Brain_Area == "Cereb" ~ "Salience/Spatial navigation",
    Brain_Area == "SN" ~ "Threat response",
    Brain_Area == "VN" ~ "Salience/Spatial navigation",
    Brain_Area == "mPFC" ~ "Cortico-striatal",
    Brain_Area == "CPu" ~ "Cortico-striatal",
    Brain_Area == "NAc" ~ "Cortico-striatal",
    Brain_Area == "M1" ~ "Cortico-striatal",
    Brain_Area == "Hypoth" ~ "Threat response",
    Brain_Area == "Thal" ~ "Salience/Spatial navigation",
    Brain_Area == "Amyg" ~ "Threat response",
    Brain_Area == "Amyg" ~ "Threat response",
    Brain_Area == "CA3" ~ "Salience/Spatial navigation",
    Brain_Area == "V1" ~ "Cortico-striatal"),.after = StructureAcronym) %>%
    filter(Brain_Area %in% c("Thal", "PAG", "VN", "Cereb", "Hypoth", "DG", 
                                "CA3", "Amyg", "CPu", "NAc", "mPFC", "mOFC",    
                                "M1","V1", "SN", "VTA"))
## Table with Brain-areas, sub-areas and network
knitr::kable(mainAreas %>% dplyr::select(Brain_Area, StructureAcronym, Group) %>%
               unique() %>% rename(`Main-Area` = Brain_Area, 
                                   `Sub-Area` = StructureAcronym, 
                                   Network = Group) %>%
               arrange(`Main-Area`), 
             caption = "Allen brain atlas main- and sub-areas") %>% 
  kableExtra::kable_styling(full_width = F) %>%
  kableExtra::scroll_box(width = "500px", height = "400px")
```

Allen brain atlas main- and sub-areas

| Main-Area | Sub-Area | Network |
| --- | --- | --- |
| Amyg | BLP | Threat response |
| Amyg | BLAp | Threat response |
| Amyg | BLA | Threat response |
| Amyg | BLAa | Threat response |
| Amyg | BMA | Threat response |
| Amyg | BMAp | Threat response |
| Amyg | BMP | Threat response |
| Amyg | BLAv | Threat response |
| Amyg | BMAa | Threat response |
| CA3 | CA3slm | Salience/Spatial navigation |
| CA3 | CA3 | Salience/Spatial navigation |
| CA3 | CA3sp | Salience/Spatial navigation |
| CA3 | CA3sr | Salience/Spatial navigation |
| CA3 | CA3slu | Salience/Spatial navigation |
| CA3 | CA3so | Salience/Spatial navigation |
| CPu | STRd | Cortico-striatal |
| CPu | CP | Cortico-striatal |
| CPu | Cau | Cortico-striatal |
| Cereb | ANcr1 | Salience/Spatial navigation |
| Cereb | ANcr1mo | Salience/Spatial navigation |
| Cereb | ANcr1gr | Salience/Spatial navigation |
| Cereb | CB | Salience/Spatial navigation |
| DG | DG-mo | Salience/Spatial navigation |
| DG | DG | Salience/Spatial navigation |
| DG | DG-sg | Salience/Spatial navigation |
| DG | DGs | Salience/Spatial navigation |
| DG | DGMol | Salience/Spatial navigation |
| DG | DGGran | Salience/Spatial navigation |
| DG | DG-po | Salience/Spatial navigation |
| DG | DGi | Salience/Spatial navigation |
| DG | DGHil | Salience/Spatial navigation |
| Hypoth | PVHpm | Threat response |
| Hypoth | PVHpml | Threat response |
| Hypoth | PVHm | Threat response |
| Hypoth | PVHmm | Threat response |
| Hypoth | PVHmpd | Threat response |
| Hypoth | PVH | Threat response |
| Hypoth | PVHp | Threat response |
| Hypoth | PVHap | Threat response |
| Hypoth | PVHlp | Threat response |
| Hypoth | PVHd | Threat response |
| M1 | MOp6a | Cortico-striatal |
| M1 | MOp6b | Cortico-striatal |
| M1 | MOp5 | Cortico-striatal |
| M1 | MOp | Cortico-striatal |
| M1 | MOp1 | Cortico-striatal |
| M1 | MOp2/3 | Cortico-striatal |
| NAc | VStr | Cortico-striatal |
| NAc | AcbSh | Cortico-striatal |
| NAc | AcbCo | Cortico-striatal |
| PAG | p1Lim-PAG | Threat response |
| PAG | TGDL-PAG | Threat response |
| PAG | TGL-PAG | Threat response |
| PAG | SCL-PAG | Threat response |
| PAG | SCDL-PAG | Threat response |
| PAG | ICDL-PAG | Threat response |
| PAG | PIsDL-PAG | Threat response |
| PAG | PIsL-PAG | Threat response |
| PAG | PB-PAG | Threat response |
| PAG | isLim-PAG | Threat response |
| PAG | PAG | Threat response |
| PAG | m1Lim-PAG | Threat response |
| PAG | Ist-PAG | Threat response |
| PAG | m1B-PAG | Threat response |
| PAG | p1B-PAG | Threat response |
| PAG | PcPL-PAG | Threat response |
| PAG | JcPL-PAG | Threat response |
| PAG | PcPV-PAG | Threat response |
| PAG | JcPV-PAG | Threat response |
| PAG | CoPV-PAG | Threat response |
| PAG | m1AD-PAG | Threat response |
| PAG | PIsD-PAG | Threat response |
| SN | SNr | Threat response |
| SN | SNc | Threat response |
| Thal | CL | Salience/Spatial navigation |
| Thal | CM | Salience/Spatial navigation |
| Thal | PVT | Salience/Spatial navigation |
| Thal | MD | Salience/Spatial navigation |
| Thal | MDc | Salience/Spatial navigation |
| Thal | MDl | Salience/Spatial navigation |
| Thal | MED | Salience/Spatial navigation |
| Thal | MDm | Salience/Spatial navigation |
| Thal | ILM | Salience/Spatial navigation |
| V1 | VISp | Cortico-striatal |
| V1 | VISp4 | Cortico-striatal |
| V1 | VISp1 | Cortico-striatal |
| V1 | VISp2/3 | Cortico-striatal |
| V1 | VISp6b | Cortico-striatal |
| V1 | VISp6a | Cortico-striatal |
| VN | MV | Salience/Spatial navigation |
| VN | SUV | Salience/Spatial navigation |
| VN | SPIV | Salience/Spatial navigation |
| VN | VNC | Salience/Spatial navigation |
| VN | LAV | Salience/Spatial navigation |
| VTA | VTA | Salience/Spatial navigation |
| mOFC | ORBm6a | Cortico-striatal |
| mOFC | ORBm2 | Cortico-striatal |
| mOFC | ORBm1 | Cortico-striatal |
| mOFC | ORBm2/3 | Cortico-striatal |
| mOFC | ORBm | Cortico-striatal |
| mOFC | ORBm5 | Cortico-striatal |
| mOFC | ORB | Cortico-striatal |
| mPFC | ACAd5 | Cortico-striatal |
| mPFC | ACAd2/3 | Cortico-striatal |
| mPFC | ACAd | Cortico-striatal |
| mPFC | ACA | Cortico-striatal |
| mPFC | ACAd1 | Cortico-striatal |
| mPFC | ACAv2/3 | Cortico-striatal |
| mPFC | ACAv | Cortico-striatal |
| mPFC | ACAv5 | Cortico-striatal |
| mPFC | ACAv1 | Cortico-striatal |
| mPFC | PL1 | Cortico-striatal |
| mPFC | PL2 | Cortico-striatal |
| mPFC | PL2/3 | Cortico-striatal |
| mPFC | PL | Cortico-striatal |
| mPFC | PL5 | Cortico-striatal |
| mPFC | ACAd6a | Cortico-striatal |
| mPFC | ACAv6a | Cortico-striatal |
| mPFC | ACAv6b | Cortico-striatal |
| mPFC | ACAd6b | Cortico-striatal |
| mPFC | CCx | Cortico-striatal |
| mPFC | ILA6a | Cortico-striatal |
| mPFC | PL6a | Cortico-striatal |
| mPFC | ILA6b | Cortico-striatal |
| mPFC | PL6b | Cortico-striatal |
| mPFC | ILA1 | Cortico-striatal |
| mPFC | ILA | Cortico-striatal |
| mPFC | ILA2/3 | Cortico-striatal |
| mPFC | ILA5 | Cortico-striatal |
| mPFC | ILA2 | Cortico-striatal |

```
## Average gene expression per brain area
mainAreas <- mainAreas %>%
    dplyr::rename(HumanGeneID = ID) %>%
    dplyr::mutate(exprs = as.numeric(exprs)) %>%
    group_by(Brain_Area, HumanGeneID) %>%
    mutate(mainsub_exprs = mean(exprs, na.omit = T)) %>% 
    dplyr::select(-c('exprs', 'Structure', 'StructureAcronym')) %>%
    rename(exprs = mainsub_exprs) %>%
    unique() %>%
    pivot_wider(names_from = "HumanGeneID", values_from = "exprs")
```

Finally, the dataset was filtered for the 946 identified
mitochondrial genes:

```
## Read mitocarta
gene_to_ID_mitocarta_hm <- readxl::read_xls(here::here("Data", "HumanMitoCarta3_0.xls"), sheet = 2) %>%
  dplyr::select(HumanGeneID,  Symbol) %>%
  unique() %>%
  mutate(HumanGeneID = as.character(HumanGeneID)) 
## Get gene IDs and symbols
mitoIDs_hm <- unique(gene_to_ID_mitocarta_hm$HumanGeneID)
mitoGenes_hm <- unique(gene_to_ID_mitocarta_hm$Symbol)
## Filter for mito gene IDs in the allen dataset
mainAreas_mito <- mainAreas %>%
  pivot_longer(cols = -c( "Brain_Area", "Group")) %>%
  dplyr::filter(name %in% mitoIDs_hm) %>%
  dplyr::mutate(value = as.numeric(value)) %>%
  pivot_wider(names_from = "name", values_from = "value")
```

##### Figure 4b - Principal component analysis

The mitochondrial gene expression values for each of the 16
main-areas were projected on a 2D and 3D principal component analysis
(PCA), and the gene contributions to PC1, PC2, and PC3 are displayed
from strongest (left) to weakest (right).

**2D PCA:**

```
## Create a new dataframe with gene symbols instead of IDs (for the pc contributions)
mainAreas_mito_pca <- mainAreas_mito %>%
  pivot_longer(cols = -c(Group, Brain_Area), names_to = "HumanGeneID") %>%
  full_join(gene_to_ID_mitocarta_hm, by= "HumanGeneID") %>%
  na.omit() %>%
  dplyr::select(-HumanGeneID) %>%
  pivot_wider(names_from = "Symbol", values_from = "value")
pca <- prcomp(mainAreas_mito_pca[,-c(1:2)], scale. = T)
top <- pca$rotation
summary_pca <- summary(pca)
p <- autoplot(pca, data = mainAreas_mito_pca,colour = 'Brain_Area', 
              size = 3)+
  theme_bw()+
  # Network color code:
  scale_color_manual(values = c("mOFC" = "#EB539F", 
                                "VTA" = "#B260EA",
                                "DG"= "#B260EA",
                                "PAG"= "#2032F5",
                                "Cereb"= "#B260EA",
                                "SN" = "#2032F5",
                                "VN"= "#B260EA",
                                "mPFC"= "#EB539F",
                                "CPu"= "#EB539F",
                                "NAc"= "#EB539F",
                                "M1"= "#EB539F",
                                "Hypoth"= "#2032F5",
                                "Thal"= "#B260EA",
                                "Amyg"= "#2032F5",
                                "CA3"= "#B260EA",
                                "V1"= "#EB539F")) +
  theme(axis.text = element_text(size = 14),
        axis.title = element_text(size = 14, face = "bold"),
        legend.text = element_text(size = 14),
        legend.title = element_blank(),
        legend.position = "right")
plotly::ggplotly(p)
```

```
p <- autoplot(pca, data = mainAreas_mito_pca,colour = 'Brain_Area', x=2, y=3,
              size = 3)+
  theme_bw()+
  scale_color_manual(values = c("mOFC" = "#EB539F",
                                "VTA" = "#B260EA",
                                "DG"= "#B260EA",
                                "PAG"= "#2032F5",
                                "Cereb"= "#B260EA",
                                "SN" = "#2032F5",
                                "VN"= "#B260EA",
                                "mPFC"= "#EB539F",
                                "CPu"= "#EB539F",
                                "NAc"= "#EB539F",
                                "M1"= "#EB539F",
                                "Hypoth"= "#2032F5",
                                "Thal"= "#B260EA",
                                "Amyg"= "#2032F5",
                                "CA3"= "#B260EA",
                                "V1"= "#EB539F")) +
  theme(axis.text = element_text(size = 14),
        axis.title = element_text(size = 14, face = "bold"),
        legend.text = element_text(size = 14),
        legend.title = element_blank(),
        legend.position = "right")
plotly::ggplotly(p)
```

**3D PCA (figure 4b):**

```
  var_1 <- round(summary_pca$importance[2,1]*100,2)
  var_2 <- round(summary_pca$importance[2,2]*100,2)
  var_3 <- round(summary_pca$importance[2,3]*100,2)
  group_color_df <- data.frame(Color = c("mOFC" = "#EB539F",
                                "VTA" = "#B260EA",
                                "DG"= "#B260EA",
                                "PAG"= "#2032F5",
                                "Cereb"= "#B260EA",
                                "SN" = "#2032F5",
                                "VN"= "#B260EA",
                                "mPFC"= "#EB539F",
                                "CPu"= "#EB539F",
                                "NAc"= "#EB539F",
                                "M1"= "#EB539F",
                                "Hypoth"= "#2032F5",
                                "Thal"= "#B260EA",
                                "Amyg"= "#2032F5",
                                "CA3"= "#B260EA",
                                "V1"= "#EB539F")) %>%
    rownames_to_column("Group")
  df <- pca$x
  df <- data.frame(PC1=df[,1], PC2=df[,2], PC3=df[,3], 
                   Group = as.factor(mainAreas_mito_pca$Brain_Area)) %>%
    full_join(group_color_df, by = "Group") %>%
    na.omit() 
  with(df, rgl::plot3d(PC1,PC2,PC3, col= Color, alpha = 0.6, size = 8, type = "p",
                       xlab = paste0("PC1 (",var_1, "%)"),
                       ylab = paste0("PC2 (",var_2, "%)"),
                       zlab = paste0("PC3 (",var_3, "%)")))

  rglwidget()
```

**Gene contributions to PC1, PC2, and PC3:**

```
pc1 <- factoextra::fviz_contrib(pca,
                 choice = "var",
                 axes = 1,
                 color = 'grey', barfill  = 'blue4',fill ='blue4',size = 0.2,
                 title = "Contributions PC1") +
  theme_minimal() +
  theme(axis.text.x = element_blank(),
        axis.title.x = element_blank(),
        axis.ticks.length.x = element_blank(),
        panel.grid.major = element_blank(), 
        panel.grid.minor = element_blank())
plotly::ggplotly(pc1)
```

```
pc2 <- factoextra::fviz_contrib(pca,
                 choice = "var",
                 axes = 2,
                 color = 'grey', barfill  = 'blue4',fill ='blue4',size = 0.2,
                 title = "Contributions PC2") +
  theme_minimal() +
  theme(axis.text.x = element_blank(),
        axis.title.x = element_blank(),
        axis.ticks.length.x = element_blank(),
        panel.grid.major = element_blank(), 
        panel.grid.minor = element_blank())
plotly::ggplotly(pc2)
```

```
pc3 <- factoextra::fviz_contrib(pca,
                 choice = "var",
                 axes = 3,
                 color = 'grey', barfill  = 'blue4',fill ='blue4',size = 0.2,
                 title = "Contributions PC3") +
  theme_minimal() +
  theme(axis.text.x = element_blank(),
        axis.title.x = element_blank(),
        axis.ticks.length.x = element_blank(),
        panel.grid.major = element_blank(), 
        panel.grid.minor = element_blank())
plotly::ggplotly(pc3)
```

```
rm(p, pca, summary_pca, df, group_color_df, var_1, var_2, var_3, top)
```

We performed a test of robustness and sensitivity by repeating these
analyses using all microscopic sub-areas individually, color-coded by
the 16 main-areas they belong to:

```
sub_Areas  <- data_raw  %>%
  #create new dataframe with column sub-area and the main-area it belongs to:
  mutate(Area = case_when(
    StructureAcronym == "ORB" ~ "Main",
    StructureAcronym == "VTA" ~ "Main",
    StructureAcronym == "DG" ~ "Main",
    StructureAcronym == "PAG" ~ "Main",
    StructureAcronym == "CB" ~ "Main",
    StructureAcronym == "SNr" ~ "Main",
    StructureAcronym == "SNc" ~ "Main",
    StructureAcronym == "VNC" ~ "Main",
    StructureAcronym == "ILA" ~ "Main",
    StructureAcronym == "CP" ~ "Main",
    StructureAcronym == "ACB" ~ "Main",
    StructureAcronym == "MOp" ~ "Main",
    StructureAcronym == "PVH" ~ "Main",
    StructureAcronym == "Th" ~ "Main",
    StructureAcronym == "TH" ~ "Main",
    StructureAcronym == "BLA" ~ "Main",
    StructureAcronym == "BMA" ~ "Main",
    StructureAcronym == "CA3" ~ "Main",
    StructureAcronym == "VISp" ~ "Main",
    TRUE ~ "Sub")) %>%
  mutate(tokeep = case_when(
    (Area == "Sub" | StructureAcronym %in% c("VTA", "SNr", "SNc")) ~TRUE,
    TRUE ~ FALSE
  )) %>%
  ## Assign main-area to sub-area
  dplyr::filter(tokeep == TRUE) %>%
  dplyr::select(-tokeep) %>%
  dplyr::mutate(Main_Area = case_when(
    StructureAcronym %in% c("ORBm6a","ORBm2","ORBm1","ORBm2/3","ORBm","ORBm5", "ORB") 
    ~ "ORB",
    StructureAcronym %in% c("VTA")
    ~ "VTA",
    StructureAcronym %in% c("DG-mo","DG","DGMol","DG-sg","DGGran","DG-po","DGs","DGi",
                            "DGHil", "DG") 
    ~ "DG",
    StructureAcronym %in% c("PcPL-PAG", "JcPL-PAG", "PcPV-PAG", "JcPV-PAG", "CoPV-PAG", 
                            "m1AD-PAG", "PIsD-PAG", "p1Lim-PAG", "TGDL-PAG", 
                            "TGL-PAG", "SCL-PAG", "SCDL-PAG", "m1Lim-PAG","ICDL-PAG", 
                            "PIsDL-PAG", "PB-PAG", "PIsL-PAG", "isLim-PAG","m1B-PAG", 
                            "p1B-PAG", "Ist-PAG", "PAG") 
    ~ "PAG",
    StructureAcronym %in% c("ANcr1", "ANcr1gr", "ANcr1mo", "CB") 
    ~ "CB",
    StructureAcronym %in% c("MV", "LAV", "SPIV", "SUV", "VNC") ~ "VNC",
    StructureAcronym %in% c("ILA6b", "ILA", "ILA2/3", "ILA5", "ILA2", "ILA1", "ILA6a", 
                            "PL6b", "PL6a", "PL1", "PL2/3", "PL", "PL2", "PL5", "ACAd5", 
                            "ACAd2/3",  "ACAd","ACA", "ACAd1", "ACAv2/3", "ACAv", "ACAv1", 
                            "ACAv5", "ACAv6a", "ILA","ACAd6a", "ACAv6b", "ACAd6b", "CCx") 
    ~ "ILA",
    StructureAcronym %in% c("STRd", "Cau","CP") 
    ~ "CP",
    StructureAcronym %in% c("AcbSh", "AcbCo", "VStr") 
    ~ "ACB",
    StructureAcronym %in% c("MOp1", "MOp2/3", "MOp5", "MOp6b", "MOp6a", "MOp") 
    ~ "MOp",
    StructureAcronym %in% c("PVHIp", "PVHd", "PVHpm", "PVHpml", "PVHm", "PVHmm", "PVHmpd", 
                            "PVHmdp", "PVHp", "PVHap", "PVH", "PVHlp") ~ "PVH",
    StructureAcronym %in% c("CL", "CM", "MDc", "MD", "MED", "MDI", "ILM", "MDm", "PVT", 
                            "TH", "MDl") 
    ~ "TH",
    StructureAcronym %in% c("BLA", "BLAa", "BLP", "BLAp", "BLA")
    ~ "BLA",
    StructureAcronym %in% c("BMAp", "BMP", "BLAv", "BMAa", "BMA") 
    ~ "BMA",
    StructureAcronym %in% c("CA3sp", "CA3sr", "CA3slu", "CA3so", "CA3slm", "CA3")
    ~ "CA3",
    StructureAcronym %in% c("VISp4", "VISp1", "VISp2/3", "VISp6a", "VISp6b", "VISp")
    ~ "VISp",
    TRUE ~StructureAcronym
  ))%>%
  ## Match acronyms to mouse dataset
  dplyr::mutate(AcronymMain = case_when(
    Main_Area == "ORB" ~ "mOFC",
    Main_Area == "VTA" ~ "VTA",
    Main_Area == "DG" ~ "DG",
    Main_Area == "PAG" ~ "PAG",
    Main_Area == "CB" ~ "Cereb",
    Main_Area == "SNr" ~ "SN",
    Main_Area == "SNc" ~ "SN",
    Main_Area == "VNC" ~ "VN",
    Main_Area == "ILA" ~ "mPFC",
    Main_Area == "CP" ~ "Cpu",
    Main_Area == "ACB" ~ "Nac",
    Main_Area == "MOp" ~ "M1",
    Main_Area == "PVH" ~ "Hypoth",
    Main_Area == "Th" ~ "Thal",
    Main_Area == "TH" ~ "Thal",
    Main_Area == "BLA" ~ "Amyg",
    Main_Area == "BMA" ~ "Amyg",
    Main_Area == "CA3" ~ "CA3",
    Main_Area == "VISp" ~ "V1",
  )) %>%
  dplyr::mutate(StructureAcronym = case_when(
    Structure == "Dentate.gyrus" ~ "DG",
    Structure == "dentate.gyrus" ~ "dg",
    Structure == "basolateral.amygdaloid.nucleus..anterior.part" ~ "BLA_ant",
    Structure == "Basolateral.amygdalar.nucleus" ~ "BLA",
    Structure == "basomedial.amygdaloid.nucleus..anterior.part" ~ "BMA_ant",
    Structure == "Basomedial.amygdalar.nucleus" ~ "BMA",
    Structure == "Field.CA3" ~ "CA3",
    Structure == "Field.CA3.1" ~ "CA3.1",
    Structure == "Field.CA3..stratum.lacunosum.moleculare.1" ~ "CA3slm.1",
    Structure == "Field.CA3..stratum.oriens.1" ~ "CA3so.1",
    Structure == "Field.CA3..pyramidal.layer" ~ "CA3sp_l",
    Structure == "Field.CA3..stratum.radiatum.1" ~ "CA3sr.1",
    Structure == "Central.lateral.nucleus.of.the.thalamus" ~ "CL_thal",
    Structure == "Central.medial.nucleus.of.the.thalamus" ~ "CM_thal",
    Structure == "Mediodorsal.nucleus.of.thalamus" ~ "MD_thal",
    TRUE ~ StructureAcronym
  )) %>%
  ## Main-area to network
  dplyr::mutate(Group = case_when(
    AcronymMain == "mOFC" ~ "Cortico-striatal",
    AcronymMain == "VTA" ~ "Salience/Spatial navigation",
    AcronymMain == "DG" ~ "Salience/Spatial navigation",
    AcronymMain == "PAG" ~ "Threat response",
    AcronymMain == "Cereb" ~ "Salience/Spatial navigation",
    AcronymMain == "SN" ~ "Threat response",
    AcronymMain == "VN" ~ "Salience/Spatial navigation",
    AcronymMain == "mPFC" ~ "Cortico-striatal",
    AcronymMain == "Cpu" ~ "Cortico-striatal",
    AcronymMain == "Nac" ~ "Cortico-striatal",
    AcronymMain == "M1" ~ "Cortico-striatal",
    AcronymMain == "Hypoth" ~ "Threat response",
    AcronymMain == "Thal" ~ "Salience/Spatial navigation",
    AcronymMain == "Thal" ~ "Salience/Spatial navigation",
    AcronymMain == "Amyg" ~ "Threat response",
    AcronymMain == "Amyg" ~ "Threat response",
    AcronymMain == "CA3" ~ "Salience/Spatial navigation",
    AcronymMain == "V1" ~ "Cortico-striatal"),.after = AcronymMain) %>%
  dplyr::rename(HumanGeneID = ID) %>%
  dplyr::mutate(exprs = as.numeric(exprs)) %>%
  pivot_wider(names_from = "HumanGeneID", values_from = "exprs") %>%
  dplyr::select(-Area)
## Filter for mitochondrial genes
subAreas_mito <- sub_Areas %>%
  pivot_longer(cols = -c("Structure"  , "StructureAcronym" ,"Main_Area","AcronymMain", "Group")) %>%
  dplyr::filter(name %in% mitoIDs_hm)%>%
  dplyr::mutate(value = as.numeric(value)) %>%
  pivot_wider(names_from = "name", values_from = "value")
## Compute the PCA
pca <- prcomp(subAreas_mito[,-c(1:5)], scale. = T)
top <- pca$rotation
summary_pca <- summary(pca)
## 3D PCA
  var_1 <- round(summary_pca$importance[2,1]*100,2)
  var_2 <- round(summary_pca$importance[2,2]*100,2)
  var_3 <- round(summary_pca$importance[2,3]*100,2)
  group_color_df <- data.frame(Color = c("mOFC" = "#EB539F",
                                         "VTA" = "#B260EA",
                                         "DG"= "#B260EA",
                                         "PAG"= "#2032F5",
                                         "Cereb"= "#B260EA",
                                         "SN" = "#2032F5",
                                         "VN"= "#B260EA",
                                         "mPFC"= "#EB539F",
                                         "Cpu"= "#EB539F",
                                         "Nac"= "#EB539F",
                                         "M1"= "#EB539F",
                                         "Hypoth"= "#2032F5",
                                         "Thal"= "#B260EA",
                                         "Amyg"= "#2032F5",
                                         "CA3"= "#B260EA",
                                         "V1"= "#EB539F")) %>%
    rownames_to_column("Group")
  df <- pca$x
  df <- data.frame(PC1=df[,1], PC2=df[,2], PC3=df[,3], Group = as.factor(subAreas_mito$AcronymMain)) %>%
    full_join(group_color_df, by = "Group") %>%
    na.omit() 
  with(df, rgl::plot3d(PC1,PC2,PC3, col= Color, alpha = 0.6, size = 8, type = "p",
                       xlab = paste0("PC1 (",var_1, "%)"),
                       ylab = paste0("PC2 (",var_2, "%)"),
                       zlab = paste0("PC3 (",var_3, "%)")))

  rglwidget()
```

##### Figure 4c - Hierarchical clustering

To compare mitochondrial gene and pathway signatures between the 16
main brain areas, each mitochondrial gene was assigned to a
mitochondrial pathway (n=149) using MitoCarta3.0 annotations. The data
was z-score transformed with a mean of 100 and a standard deviation of
10 to allow for direct gene expression comparisons between brain
areas.

```
mainAreas_mito_z_score <- mainAreas %>%
   pivot_longer(cols = -c("Brain_Area", "Group"), 
                names_to = "HumanGeneID", values_to= "exprs") %>%
  group_by(Brain_Area) %>%
  mutate(exprs = (exprs - mean(exprs, na.omit = T))/sd(exprs, na.rm = T)) %>%
  mutate(exprs = (exprs * 10) + 100) %>%
  filter(HumanGeneID %in% mitoIDs_hm) %>%
  pivot_wider(names_from = "HumanGeneID", values_from = "exprs")
```

Raw data distribution (vertical line = average gene expression in
each brain area)

```
p <- mainAreas %>%
  pivot_longer(cols = -c("Brain_Area", "Group"), 
                names_to = "HumanGeneID", values_to= "exprs") %>%
  mutate(mean_all= mean(exprs, na.omit = T)) %>%
  group_by(Brain_Area) %>%
  mutate(mean_structure = mean(exprs, na.omit = T)) %>%
  ggplot(aes(x= exprs, color = Brain_Area)) +
  geom_vline(aes(xintercept =mean_structure, color = Brain_Area), alpha = 0.2) +
  geom_line(stat = "density") +
  xlab("Gene expression") +
  theme_bw() +
  scale_y_continuous(limits = c(0, 0.14), expand = expansion(mult = c(0, .1))) 

plotly::ggplotly(p)
```

Data distribution after z-score transform with mean of 100 (vertical
line) and standard deviation of 10

```
p <- mainAreas %>%
  pivot_longer(cols = -c("Brain_Area", "Group"), 
                names_to = "HumanGeneID", values_to= "exprs") %>%
  group_by(Brain_Area) %>%
  mutate(exprs = (exprs - mean(exprs, na.omit = T))/sd(exprs, na.rm = T)) %>%
  mutate(exprs = (exprs * 10) + 100) %>%
  mutate(mean_all= mean(exprs, na.omit = T)) %>%
  group_by(Brain_Area) %>%
  mutate(mean_structure = mean(exprs, na.omit = T)) %>%
  ggplot(aes(x= exprs, color = Brain_Area)) +
  geom_line(stat = "density") +
  geom_vline(aes(xintercept =mean_structure, color = Brain_Area), alpha = 0.2) +
  xlab("Gene expression")+
  theme_bw()+
  scale_y_continuous(limits = c(0, 0.04), expand = expansion(mult = c(0, .1))) 

plotly::ggplotly(p)
```

From the transformed data, the expression of genes in a given pathway
(as annotated in MitoCarta3.0) were averaged, yielding 149 mitochondrial
pathway scores for each brain area.

Hierarchical clustering (Figure 4c) of the resulting matrix (16 brain
areas x 149 pathways) was performed using the Euclidean distance
calculated from relative pathway scores and the ward.D2 method.

```
pathway_score <- mainAreas_mito_z_score %>%
  pivot_longer(-c("Group","Brain_Area"), 
               names_to = "ID", values_to = "exprs") %>%
  full_join(pathway_gene_ID, by = "ID") %>%
  group_by(Pathway, Brain_Area) %>%
  mutate(Average_exprs = mean(exprs)) %>%
  ungroup() %>%
  dplyr::select( Group, Average_exprs, Pathway, Brain_Area) %>%
  unique() %>%
  na.omit() %>%
  pivot_wider(names_from = "Pathway", values_from = "Average_exprs") %>%
  column_to_rownames("Brain_Area") %>%
  mutate(Group = case_when(
    Group == "Cortico-striatal"~"Cortico-striatal",
     Group ==         "Salience/Spatial navigation"~"Salience/Spat.Nav.",
     Group ==          "Threat response" ~ "Threat response"))

exprs <-t(scale(pathway_score[,2:ncol(pathway_score)])) # scale columnwise to compare brain areas relative to each other
col_fun = colorRamp2(c(range(exprs)[1], 0, range(exprs)[2]),c("blue", "white", "red")) 
### Clustering Rows
row_dist    = dist(as.matrix(exprs), method="euclidean")
rowdend     = hclust(row_dist, method="ward.D2")
column_dist    = dist(as.matrix(t(exprs)), method="euclidean",)
columndend     = hclust(column_dist, method="ward.D2")
## Row annotation
column_anno_df <- pathway_score %>%
  as.data.frame() %>%
  dplyr::select(Group) 
## Column annotation
column_anno = columnAnnotation(
  `Tissue group`=column_anno_df$Group,
  col=list(`Tissue group`  = color_groups),
  show_annotation_name = F,
  show_legend =  T,
  simple_anno_size = unit(0.2, "cm"),
  annotation_legend_param = list(nrow=3,
  labels_gp = gpar(fontsize = 8),
              title_gp = gpar(fontsize = 8)))
## Build the heatmap
HM <- Heatmap(exprs, 
              name = "Rel. pathway score", 
              col=col_fun,
              column_title="",
              row_title="",
              row_dend_side = "right",
              row_names_side = "left",
              row_dend_width = unit(0.2, "cm"),
              show_row_dend=TRUE,
              show_row_names=T,
              show_column_names=T,
              cluster_rows =rowdend,
              cluster_columns = columndend,
              row_names_gp = grid::gpar(fontsize = 3),
              column_title_gp = grid::gpar(fontsize = 10),
              column_names_gp = grid::gpar(fontsize = 8),
              top_annotation=column_anno,
              width = unit(70, "mm"),
              heatmap_legend_param = list(
                title = "Rel. pathway score",
              labels_gp = gpar(fontsize = 8),
              title_gp = gpar(fontsize = 8))
            
)
draw(HM)
```

##### Figure 4d - Ranked pathway scores

To quantify mitotype differences between network1 and network2/3, we
calculated pathway scores for each network group respectively using the
average expression of all genes annotated to each pathway.

```
pathway_score <- mainAreas_mito_z_score %>%
  pivot_longer(-c("Group", "Brain_Area"), 
               names_to = "ID", values_to = "exprs") %>%
  full_join(pathway_gene_ID, by = "ID") %>%
  mutate(Network = case_when(
    Group =="Cortico-striatal" ~ "Network1",
    (Group ==  "Salience/Spatial navigation" | 
       Group == "Threat response" ) ~ "Network2_3",
  )) %>%
  mutate(Group = case_when(
    Group == "Cortico-striatal"~"Cortico-striatal",
     Group ==         "Salience/Spatial navigation"~"Salience/Spat.Nav.",
     Group ==          "Threat response" ~ "Threat response")) %>%

  group_by(Pathway,Network) %>%
  mutate(Average_exprs = mean(exprs)) %>%
   ungroup() %>%
    na.omit() %>%
  dplyr::select(Network, Average_exprs, Pathway) %>%
  unique() %>%
  pivot_wider(names_from = "Network", values_from = "Average_exprs")
```

Next, we calculated the log2 fold change (log2(network1 /
network2.3)), and ranked the fold changes from lowest (higher in network
2/3) to highest (higher in network1):

```
pathway_score <- pathway_score %>%
  column_to_rownames("Pathway") %>%
  dplyr::mutate(log2FC = log2(Network1/Network2_3)) %>%
  arrange(log2FC) %>% #arrange data ascending
  rownames_to_column("Pathway_Level3") %>% 
  dplyr::mutate(xaxis = seq(1:149)) %>% #add x-axis to sort ascending from left to right
  dplyr::select(Pathway_Level3,log2FC, xaxis)

p <- pathway_score %>% 
  ggplot(aes(x =xaxis, y = log2FC, label = Pathway_Level3)) +
  geom_point(alpha = 0.7, size = 2.5, shape = 21, color = "darkgray", fill = "gray") +
 geom_hline(yintercept = 0, linetype = "dotted", color = "gray", linewidth = 0.5) +
  labs(y = "Log2 fold change Network1 to Network2_3", x = "Ranked Mitopathway scores") +
  theme_bw() +
  geom_vline(aes(xintercept =119), linetype = "dotted", color = "gray", linewidth = 0.5) +
  theme(axis.text.x = element_blank(),
        axis.title.x = element_text(size=10),
        legend.position = "none",
        axis.text.y = element_text(size = 8),
        axis.title.y = element_text(size=10),
        panel.grid.major = element_blank(), 
        panel.grid.minor = element_blank())

plotly::ggplotly(p)
```

##### Figure 4e - Bivariate plot of Vitamin B2 metabolism & G3P shuttle

From the ranked mitopathway scores we picked the top (G3P shuttle)
and bottom (Vit B2 metabolism) mitopathway and calculated the pathway
score for each brain area (as shown in figure 4c)

```
pathway_score <- mainAreas_mito_z_score %>%
  pivot_longer(-c("Group",  "Brain_Area"), 
               names_to = "ID", values_to = "exprs") %>%
  full_join(pathway_gene_ID, by = "ID") %>%
   mutate(Group = case_when(
    Group == "Cortico-striatal"~"Cortico-striatal",
     Group ==         "Salience/Spatial navigation"~"Salience/Spat.Nav.",
     Group ==          "Threat response" ~ "Threat response")) %>%

  group_by(Pathway,Brain_Area) %>%
  mutate(Average_exprs = mean(exprs)) %>%
   ungroup() %>%
    na.omit() %>%
  dplyr::select(-c(ID, exprs, Gene) )%>%
  unique() %>%
  pivot_wider(names_from = "Pathway", values_from = "Average_exprs")
```

Next, we plotted both pathways against one another, color-coded by
network:

```
color_structure <- c(
    "mOFC" = "#EB539F",
    "VTA" = "#B260EA",
    "DG" = "#B260EA",
    "PAG" = "#2032F5",
    "Cereb" = "#B260EA",
    "SN" = "#2032F5",
    "VN" = "#B260EA",
    "mPFC" = "#EB539F",
    "CPu" = "#EB539F",
    "NAc" = "#EB539F",
    "M1" = "#EB539F",
    "Hypoth" = "#2032F5",
    "Thal" = "#B260EA",
    "Amyg" = "#2032F5",
    "CA3" = "#B260EA",
    "V1" = "#EB539F")

p <- pathway_score %>%
   ggplot(aes(x = `Glycerol phosphate shuttle`, y = `Vitamin B2 metabolism`, color = Brain_Area)) +
  geom_point(alpha = 0.6, size = 4) +
  scale_color_manual(values = color_structure) +
  theme_bw() +
  theme(
    axis.title = element_text(size = 10), 
    axis.text = element_text(size =8)
  ) 
plotly::ggplotly(p)
```

##### Figure 4f - Ratio G3P/Vit. B2

We calculated the ratio of both pathways (G3P shuttle/Vitamine B2
metabolism) for each brain area individually, and plotted the ratios
from highest to lowest. We futher calculated the percent difference
between the brain area with the highest ratio and the one with the
lowest ratio:

```
ratio <- pathway_score %>%
  mutate(ratio = `Glycerol phosphate shuttle`/`Vitamin B2 metabolism`) %>%
  dplyr::select(Brain_Area, ratio, 
                `Vitamin B2 metabolism`,`Glycerol phosphate shuttle` ) %>%
    unique() %>%
  arrange(desc(ratio))

ratio$Brain_Area  <- factor(ratio$Brain_Area, levels = ratio$Brain_Area)
  p <- ratio %>% 
  ggplot(aes(x= Brain_Area, color = Brain_Area)) +
  geom_segment(aes(xend=Brain_Area,
                   y=1, yend=ratio), size=4, alpha =0.6, show.legend = F) +
  scale_color_manual(values = color_structure) +
  scale_y_continuous(limits = c(1,1.18), expand = expansion(mult = c(0, .1))) +
  labs(y= "G3P shuttle / Vit. B2 metabolism") +
  theme_classic() +
  theme( axis.title.x = element_blank(),
      axis.text.x = element_text(angle = 45, hjust = 1),
        axis.text.y = element_text(size = 8),
        axis.title.y = element_text(size = 10),
        axis.ticks.x = element_blank(),
        panel.grid.major = element_blank(), 
        panel.grid.minor = element_blank(),
        legend.position = "none") 
plotly::ggplotly(p, tooltip="ratio")
```

```
change <- ratio %>% 
  mutate(min = min(ratio), max= max(ratio)) %>%
  filter(min == ratio | max == ratio) %>%
    column_to_rownames("Brain_Area") %>%
  dplyr::select(ratio) %>%
  t()
paste("Percent difference highest to lowest:", ((change[,1] - change[,2]) / change[1] ) *100)
```

```
## [1] "Percent difference highest to lowest: 13.46755774427"
```

##### Figure 4g - Bivariate plot of Calcium homeostasis & Metabolism

```
p <- pathway_score %>%
   ggplot(aes(x = `Metabolism`, y = `Calcium homeostasis`, color = Brain_Area)) +
  geom_point(alpha = 0.6, size = 4) +
  scale_color_manual(values = color_structure) +
  theme_bw() +
  theme(
    axis.title = element_text(size = 10), 
    axis.text = element_text(size =8)
  ) 
plotly::ggplotly(p)
```

```
change <- ratio %>% 
  mutate(min = min(ratio), max= max(ratio)) %>%
  filter(min == ratio | max == ratio) %>%
    column_to_rownames("Brain_Area") %>%
  dplyr::select(ratio) %>%
  t()
```

##### Figure 4h - Ratio Calcium homeostasis/Metabolism

```
ratio <- pathway_score %>%
  mutate(ratio = `Calcium homeostasis`/`Metabolism`) %>%
  dplyr::select(Brain_Area, ratio, 
                `Metabolism`,`Calcium homeostasis` ) %>%
    unique() %>%
  arrange(desc(ratio))

ratio$Brain_Area  <- factor(ratio$Brain_Area, levels = ratio$Brain_Area)
  p <- ratio %>% 
  ggplot(aes(x= Brain_Area, color = Brain_Area)) +
  geom_segment(aes(xend=Brain_Area,
                   y=1, yend=ratio), size=4, alpha =0.6, show.legend = F) +
  scale_color_manual(values = color_structure) +
  scale_y_continuous(limits = c(1,1.04), expand = expansion(mult = c(0, .1))) +
  labs(y= "Calcium homeostasis / Metabolism") +
  theme_classic() +
  theme(axis.title.x = element_blank(),
        axis.text.x = element_text(angle = 45, hjust = 1),
        axis.text.y = element_text(size = 8),
        axis.title.y = element_text(size = 10),
        axis.ticks.x = element_blank(),
        panel.grid.major = element_blank(), 
        panel.grid.minor = element_blank(),
        legend.position = "none") 
plotly::ggplotly(p, tooltip="ratio")
```

```
change <- ratio %>% 
  mutate(min = min(ratio), max= max(ratio)) %>%
  filter(min == ratio | max == ratio) %>%
    column_to_rownames("Brain_Area") %>%
  dplyr::select(ratio) %>%
  t()
paste("Percent difference highest to lowest:", ((change[,1] - change[,2]) / change[1] ) *100)
```

```
## [1] "Percent difference highest to lowest: 1.80428841082514"
```
